# Basal stem cell fate specification is mediated by SMAD signaling in the developing human lung

**DOI:** 10.1101/461103

**Authors:** Alyssa J. Miller, Qianhui Yu, Michael Czerwinski, Yu-Hwai Tsai, Renee F. Conway, Angeline Wu, Emily M. Holloway, Taylor Walker, Ian A. Glass, Barbara Treutlein, J. Gray Camp, Jason R. Spence

**Affiliations:** Program in Cell and Molecular Biology, University of Michigan Medical School, Ann Arbor, Michigan; Department of Internal Medicine, Gastroenterology, University of Michigan Medical School, Ann Arbor, Michigan; Max Planck Institute for Evolutionary Anthropology, Leipzig, Germany; Center for Organogenesis, University of Michigan Medical School, Ann Arbor, Michigan; Department of Cell and Developmental Biology, University of Michigan Medical School, Ann Arbor, Michigan; Department of Pediatrics, Genetic Medicine, University of Washington, Seattle, Washington; Max Planck Institute of Molecular Cell Biology and Genetics, Dresden, Germany; Department of Biosciences, Technical University Munich, Freising, Germany; Department of Biomedical Engineering, University of Michigan College of Engineering, Ann Arbor, Michigan

## Abstract

Basal stem cells (basal cells), located in the bronchi and trachea of the human lung epithelium, play a critical role in normal airway homeostasis and repair, and have been implicated in the development of diseases such as cancer^1-4^. Additionally, basal-like cells contribute to alveolar regeneration and fibrosis following severe injury^5-8^. However, the developmental origin of basal cells in humans is unclear. Previous work has shown that specialized progenitor cells exist at the tips of epithelial tubes during lung branching morphogenesis, and in mice, give rise to all alveolar and airway lineages^9,10^. These ‘bud tip progenitor cells’ have also been described in the developing human lung^11-13^, but the mechanisms controlling bud tip differentiation into specific cell lineages, including basal cells, are unknown. Here, we interrogated the bud tip-to-basal cell transition using human tissue specimens, bud tip progenitor organoid cultures^11^, and single-cell transcriptomics. We used single-cell mRNA sequencing (scRNAseq) of developing human lung specimens from 15-21 weeks gestation to identify molecular signatures and cell states in the developing human airway epithelium. We then inferred differentiation trajectories during bud tip-to-airway differentiation, which revealed a previously undescribed transitional cell state (‘hub progenitors’) and implicated SMAD signaling as a regulator of the bud tip-to-basal cell transition. We used bud tip progenitor organoids to show that TGFT1 and BMP4 mediated SMAD signaling robustly induced the transition into functional basal-like cells, and these *in vitro*-derived basal cells exhibited clonal expansion, self-renewal and multilineage differentiation. This work provides a framework for deducing and validating key regulators of cell fate decisions using single cell transcriptomics and human organoid models. Further, the identification of SMAD signaling as a critical regulator of newly born basal cells in the lung may have implications for regenerative medicine, basal cell development in other organs, and understanding basal cell misregulation in disease.

## Introduction

Basal stem cells serve as epithelial stem cells in multiple organ systems, including lung, skin, esophagus, breast and prostate, where they contribute to organ homeostasis and repair after injury^1,14-20^ and their misregulation has been implicated in diseases, including cancer^4,21,22^. In the lung, basal stem cells are marked by the transcription factor TP63, which is essential for basal cell fate determination and function^23,24^, and reside on the basolateral surface of the pseudostratified epithelium of proximal airways in both mice and humans^1,2^. Resident airway basal cells routinely give rise to secretory and multiciliated cells of the airway during homeostasis and repair. In addition to basal stem cells lining the airway epithelium, basal-like cells appear in the alveolar regions of the lung after severe injury in mice. These basal-like cells express KRT5 and TP63 and contribute to injury repair^5-8^, and TP63+ basal-like cells have been identified in the distal airways of human patients with pulmonary fibrosis, suggesting these distal basal-like cells may contribute to repair and disease in humans^21^. However, the mechanisms governing basal cell fate specification during development or during disease remain unclear. Lineage tracing experiments in mice have shown that bud tip progenitor cells give rise to all lung epithelial cell types during development, including basal cells^9,10^. Similarly, human fetal bud tip progenitor cells are able to give rise to putative TP63+ basal stem cells upon transplantation *in vivo*^1-4,12^; however, our understanding of basal cell specification during human development has been hindered by the inherent challenges of performing mechanistic studies in human tissue and by limited tissue availability. Here, we interrogated the developmental mechanisms controlling the bud tip-to basal cell transition during human lung development using scRNAseq, *in situ* mRNA and protein analysis of human fetal lung tissue, and functional *in vitro* experiments utilizing human fetal-derived bud tip progenitor organoids.

## Results and discussion

### Single-cell transcriptomics defines bud tip progenitor and basal stem cell signatures in the developing human lung

We used single-cell transcriptomics to identify the epithelial cell states that arise during human lung development. We performed scRNAseq on dissociated whole distal lung, small airway, and scraped tracheal epithelial cells (Fig. 1a, Extended Data Fig. 1a) from human fetal lung tissue ranging from 15 to 21 weeks gestation, and identified diverse immune, mesenchymal, and epithelial populations (Extended Data Fig. 1b, c). We computationally extracted 7,779 epithelial cells based on clusters with high *EPCAM* expression (Extended Data Fig. 1c), then re-clustered these cells using hierarchical clustering, and visualized the heterogeneity using t-distributed Stochastic Neighbor Embedding (tSNE) (Fig. 1b). Based on this analysis, we identified 14 epithelial cell populations, and cluster identities were assigned based on known markers, where possible (Fig. 1c, d, h; Extended Data Figure 2a, Extended Data Table 1). Briefly, we identified multiple known groups of lung epithelial progenitors including bud tip progenitors (cluster 9), basal cells (cluster 12) and differentiated cell types including multiciliated cells (cluster 1), neuroendocrine cells (cluster 7), and club-like and goblet-like secretory cells (clusters 5 and 6, respectively). We also identified a novel population of epithelial progenitor cells which we term ‘hub progenitor cells’ (cluster 10). These cells express high levels of *SCGB3A2* and *SFTPB* and are molecularly distinct from other secretory cell populations; for example, they do not express high levels of the club cell marker *SCGB1A1.* Additional analysis of this population is provided in Figure 2.

**Figure 1.**
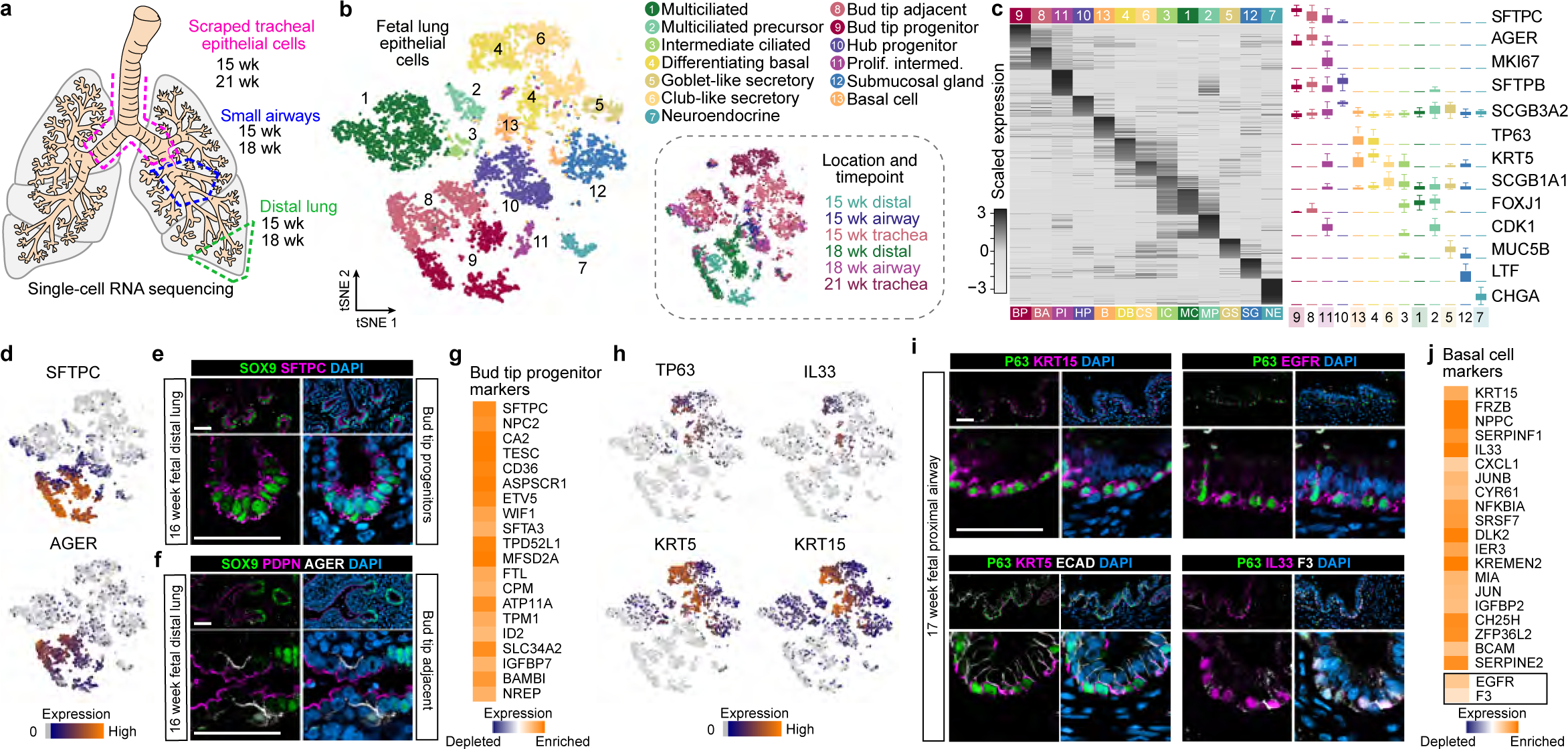
Defining cell signatures of human fetal lung epithelial cells. **a**) Schematic of the experimental setup and samples included in the analysis. **b**) A total of 7,779 EPCAM+ cells were isolated and cells were visualized using a tSNE plot revealing 13 clusters of cells. **c**) Clusters were identified by expression of markers canonically associated with cell populations based on published literature, or given a new name (e.g. hub progenitors). **d**) Feature plots show that cluster 9 contains high levels of expression of canonical bud tip progenitor marker *SFTPC* and cluster 8 contains cells that express low levels of bud tip progenitor marker *SFTPC* and also express *AGER,* a canonical marker of alveolar epithelial type 1 cells. **e**) Protein staining of a 16 week fetal lung specimen shows that bud tip progenitors stain positive for SOX9 (green) and SFTPC (pink). Scale bars represent 50 µm. **f**) Protein staining of the cells immediately adjacent to the bud tip progenitors reveals a population that is SOX9 negative (green) but which express PDPN (pink) and AGER (white). Scale bars represent 50 µm. **g**) Genes highly enriched in the bud tip progenitor cluster relative to all other lung epithelial cell clusters at 15-21 weeks gestation. **h**) Feature showing cluster 13 cells that express canonical basal cell markers *TP63* and *KRT5*, as well as other makers such as *KRT15* and *IL33*. **i**) Protein staining confirms that tracheal TP63+ basal cells in 17 week fetal lungs express basal cell markers KRT15 (pink), EGFR (pink), IL33 (pink), F3 (white) and PDPN (pink), markers that were strongly expressed in cluster 13 by scRNAseq. **j**) Genes highly enriched in the basal cell cluster relative to all other lung epithelial cell clusters15-21 weeks gestation. *EGFR* and *F3* were identified as highly enriched cell surface markers.

**Extended Data Figure 1.**
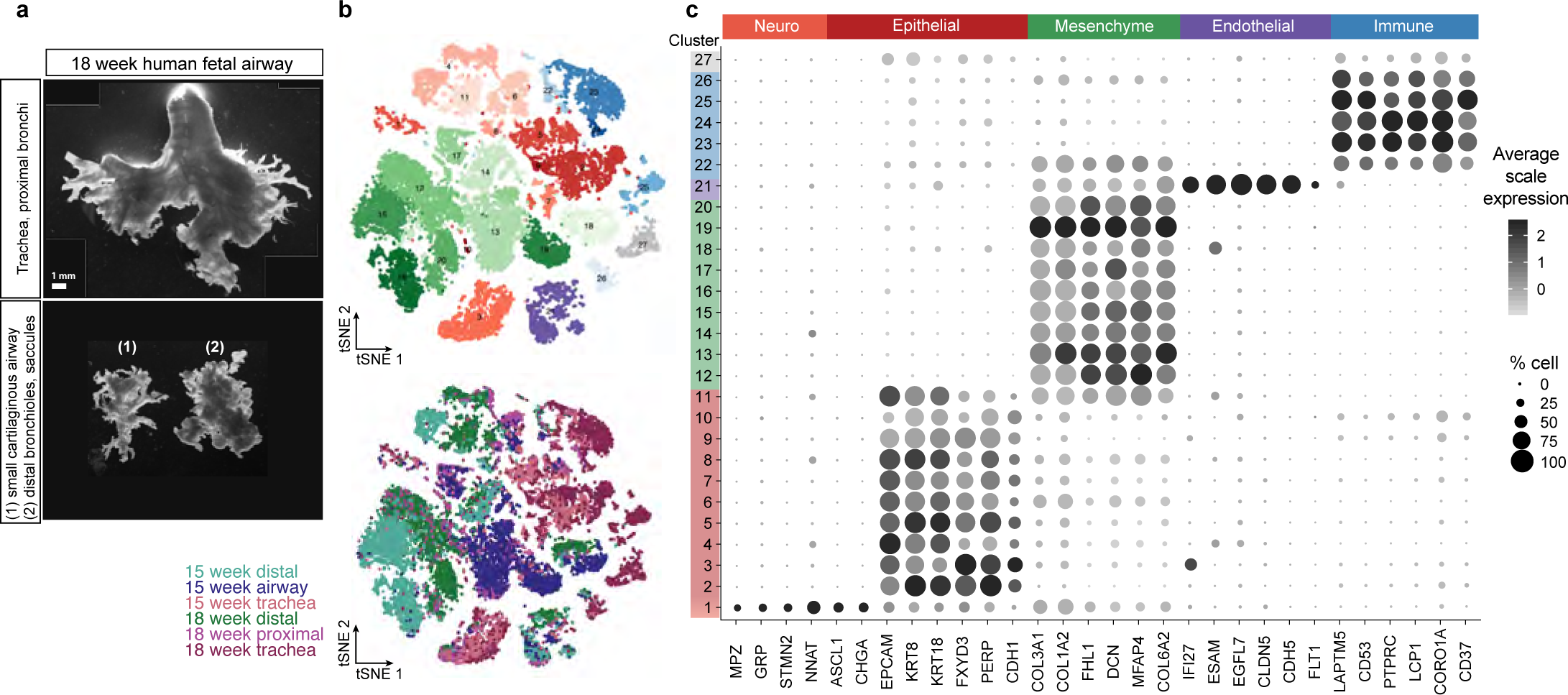
Fetal lung characterization by scRNAseq. **a**) Brightfield images of 18 week dissected human fetal lung tissue. Top panel shows the trachea and main bronchi. To collect cells for scRNAseq from the trachea, the tracheal tube was cut lengthwise, and epithelial cells were scraped with a scalpel and processed for scRNAseq. Bottom panel shows the small cartilaginous airways (1) and the distal lung (2). Pieces of tissue this size were homogenized in full to single cells and subjected to scRNAseq. For all groups, EPCAM+ cells were later isolated from the dataset for downstream analysis. Scale bar represents 1mm. **b**) Heirarchical clustering was performed on all cells from scRNAseq and visualized by tSNE. This analysis revealed 27 clusters. **d**) Cell clusters were identified based on expression of known and canonical marker genes for cell types, including those of the neural, epithelial, mesenchymal, endothelial, and immune populations. Cells with high EPCAM expression (clusters 1-11) were extracted for downstream analysis.

**Figure 2.**
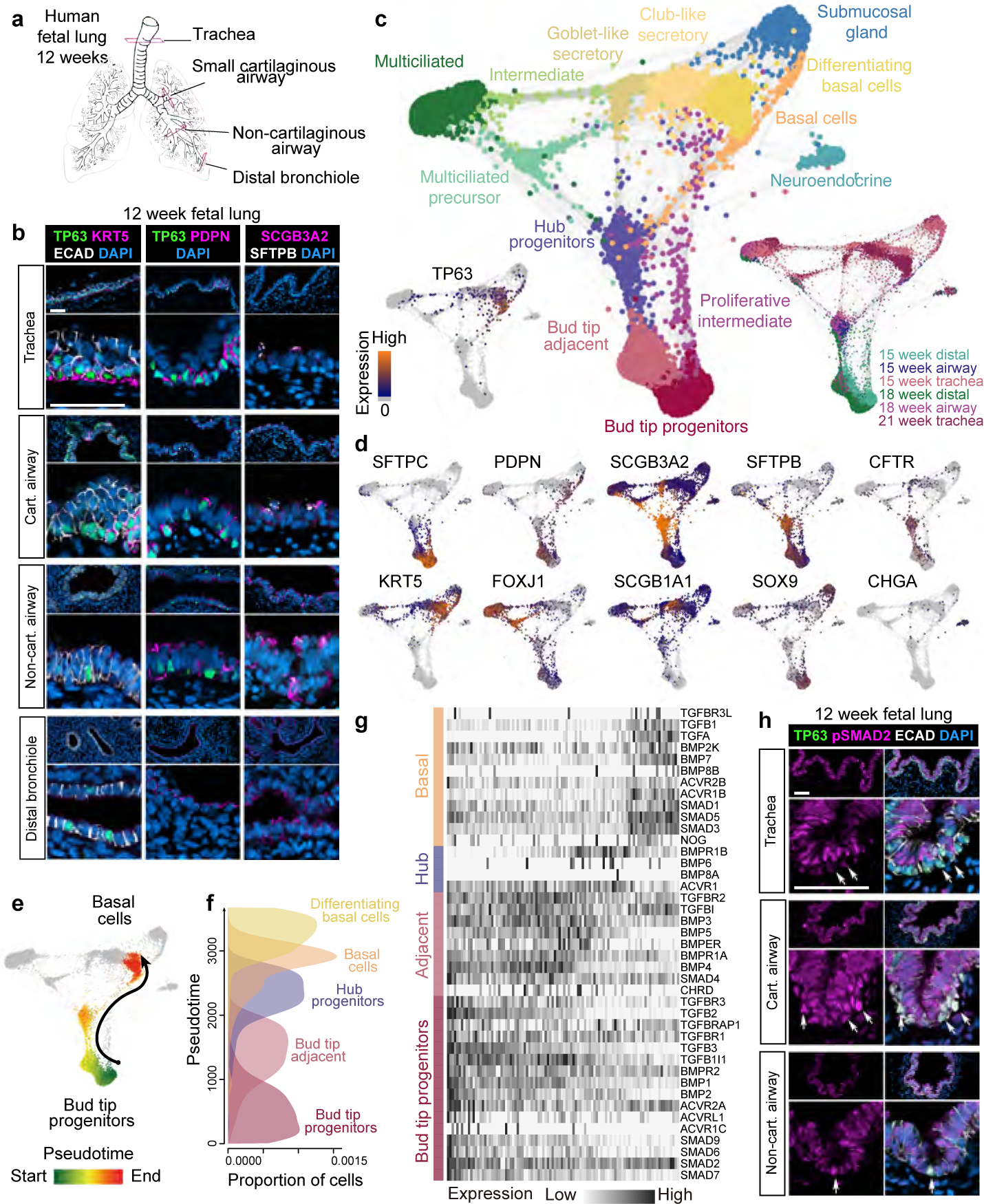
A distal-to-proximal cellular hierarchy identifies a novel intermediate ‘hub progenitor’ cell. **a**) Schematic of a 12 week human fetal lung showing the areas of analysis along the proximal-distal axis of the airway epithelium. **b**) Protein staining of 12 week fetal lungs along the proximal-distal axis for TP63 (green) and KRT5 (pink), left panel; co-staining of TP63 (green) with PDPN (pink), middle panel; co-staining for hub cell markers SCGB3A2 and SFTPB, right panel. Representative images shown from n=3 independent specimens. Scale bars represent 50 µm. **c**) k-nearest neighbor network construction was performed and visualized with SPRING^27^ (center plot), with cell identifications based on cluster IDs from Fig. 1. Locations of cells from each specimen are shown on the SPRING plot (right), and *TP63*+ cells are highlighted (left). **d**) Expression of individual genes were overlaid on the SPRING plot. *SFTPC* marks the bud tip progenitor cluster, *PDPN* marks the bud tip adjacent/newly differentiating bud tip progenitor population. Hub progenitors express high levels of *SCGB3A2* and *SFTPB.* Markers of more differentiated cells, including *KRT5* (basal cells), *FOXJ1* (ciliated cells), *SCGB1A1* (secretory/club cells) and *CHGA* (neuroendocrine cells) identify the remaining clusters. **e**) Cells appear to have continuous gene expression topologies from bud tip progenitor to the basal cell lineage. **f**) These cells were extracted, and the rank of diffusion component 1 was then used to determine pseudotime, which identified a trajectory from bud tip progenitors to bud tip adjacent to hub progenitor to basal cell. Pseudotime distribution shows the proportion of cells in each cluster at different pseudotimes, showing that the majority of cells at early pseudotime points are bud tip progenitors, and, as pseudotime progresses, move through bud tip adjacent, hub progenitor, and basal cell populations. **g**) Expression of TGF) pathway associated genes across pseudotime shows variable expression of certain signaling elements from bud tip progenitors (dark green) to bud tip adjacent (light green) to hub progenitors (light purple) to basal cells (dark purple). **h**) Protein staining of 12 week fetal lungs showed strong nuclear phospho-SMAD2 staining (pSMAD2; pink) in TP63+ cells (green) in the small-and non-cartilaginous airways, whereas TP63+ cells within the trachea show very low to no nuclear pSMAD2 staining. Scale bars represent 50 µm. Representative images shown from n=3 independent specimens.

We next sought to utilize the single cell transcriptomic data to identify gene signatures and to validate protein markers for bud tip progenitors and basal stem cells. We observed that cells in cluster 9 exhibited the strongest expression of canonical markers of human bud tip progenitors, including *SOX9*, *SFTPC, ETV5* and *ID2* (Fig. 1d)^5,7,11,12,25^. Protein expression of certain bud tip marker genes was confirmed by protein staining in fetal lung tissue (Fig. 1e; n=3 biological replicates). Cluster 8 contained cells co-expressing certain bud tip progenitor markers (low *SFTPC*, *ID2*) as well as canonical alveolar epithelial type 1 (AECI) markers such as *HOPX*, *PDPN* and *AGER* (Fig. 1f). Protein staining of the human fetal lung at 16 weeks shows that the cells directly adjacent to the SOX9+ bud tips do not express SOX9, but do express AECI markers PDPN and AGER, suggesting this population may be newly differentiating bud tip progenitors (Fig. 1e; n=3 biological replicates). We refer to this population as ‘bud tip adjacent’ cells in subsequent analysis. A comparison of genes expressed in the bud tip progenitor cell cluster relative to expression in all other clusters allowed us to identify marker genes most highly enriched in bud tip progenitors (cluster 9), thereby defining a transcriptional signature of human fetal bud tip progenitors at 15-18 weeks gestation (Fig. 1g, Extended Data Fig. 2b, Extended Data Table 1).

**Extended Data Figure 2.**
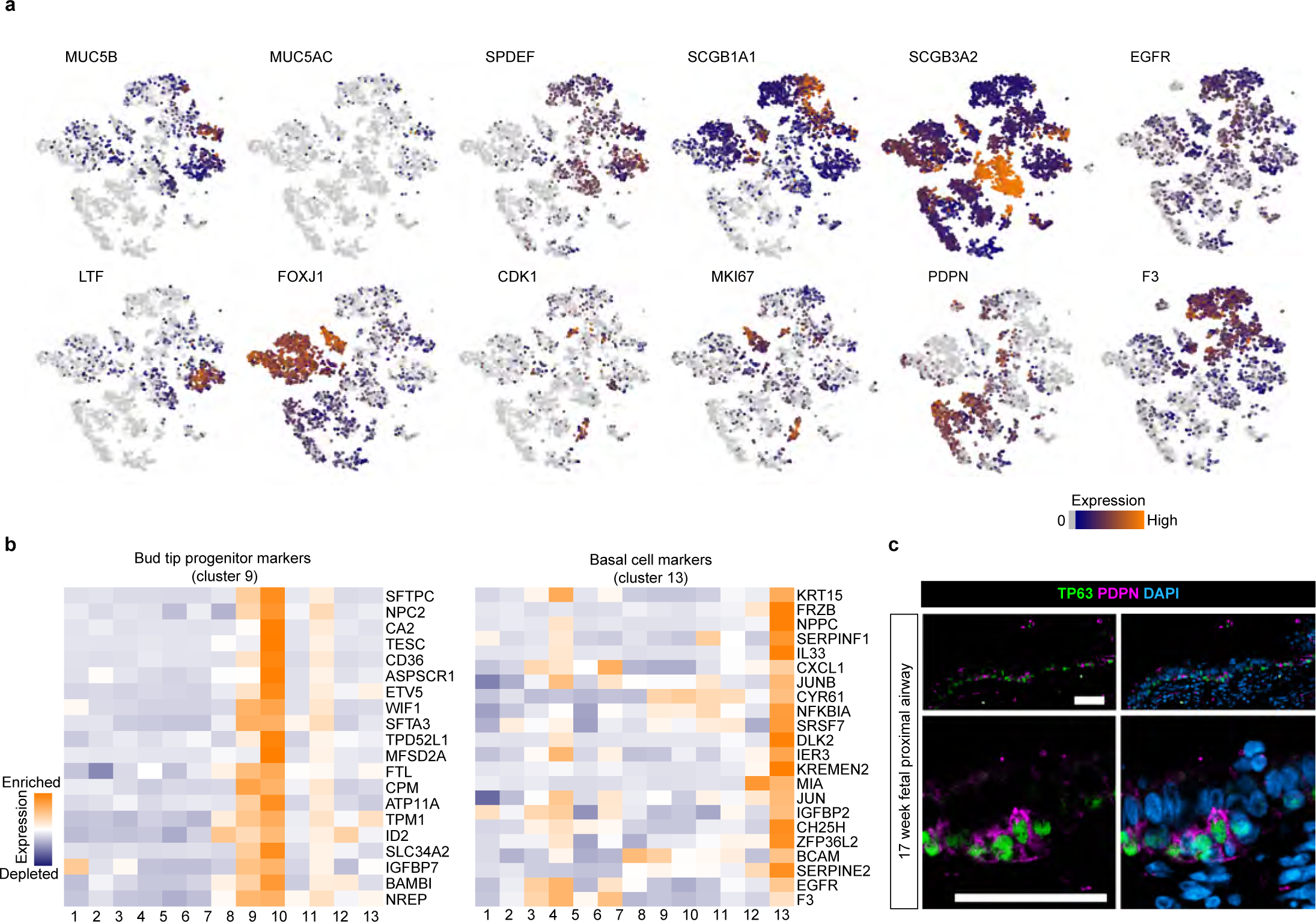
Bud tip progenitor and basal cell profiles. **a**) Feature plots of tSNE clustering show expression of markers for lung epithelial cell types, including goblet cell markers *MUC5B* and *MUC5AC*, secretory marker *SPDEF*, club cell marker *SCGB1A1,* hub progenitor marker *SCGB3A2*, submucosal gland cell marker LTF, multiciliated cell marker FOXJ1 and proliferation markers MKI67 and CDK1. **b**) Heatmaps show the 20 most enriched (orange) genes for bud tip progenitors (cluster 9) or basal cells (cluster 13) relative to all other in all epithelial clusters from 15-21 week gestation human fetal lung specimens. **c**) TP63+ basal cells (green) in the trachea of 17 week fetal lungs (n=3 biological replicates) stain positive for surface marker PDPN (pink). Scale bar represents 50 µm.

To define a basal stem cell signature, we first identified basal cells as cells within cluster 13 based on expression of canonical basal cell markers *TP63* and *KRT5* (Fig. 1c, h)^1,2,9,10,12,26^. In addition, this cluster highly expressed a set of basal cell-enriched genes that included *KRT15, IL33*, *S100A2*, *F3, EGFR*, and *PDPN* (Fig. 1h, Extended Data Fig. 2a; Extended Data Table 1), and protein staining validated that KRT5, IL33, F3, EGFR, KRT15 and PDPN are co-expressed with TP63+ cells in the trachea at 17-weeks of gestation (Fig. 1i; Extended Data Fig. 2c; n=3 biological replicates). Comparison of the average gene expression levels in basal cells (cluster 13) to the average gene expression in all other clusters allowed us to identify a list of the most enriched genes in basal cells, defining a transcriptional signature for this cell population (Fig. 1j, Extended Data Fig. 2b; Extended Data Table 1). Importantly, we manually identified EGFR and F3 as cell surface markers that are co-expressed in basal cells that could be used to isolate these cells using fluorescence activated cell sorting (FACS) techniques. Together, these data provide a reference atlas of epithelial cell states with defined molecular signatures that arise in the human developing lung epithelium.

### Continuous gene expression topologies suggest a hierarchy of cell identities along the distal to proximal axis and identifies hub progenitors as a new intermediate cell type

Next, we utilized analysis of scRNAseq data, protein staining and mRNA fluorescent *in situ* hybridization (FISH) to gain insights into cell fates and transitions in the developing human lung epithelium. Recent work in the mouse utilizing *TP63* lineage tracing identified populations of “early” basal cells that appear early in development and expressed TP63 but not KRT5 and give rise to all epithelial cell types in the mature lung, and “late” basal cells that express TP63 and KRT5 within the mouse trachea and have lineage restriction to the upper airway^10-13^. To begin to interrogate this concept of “early” and “late” basal cells in the context of basal cell lineage specification in human lungs, we performed protein staining and FISH on 12 week fetal lungs and imaged regions along the bud tip-to-trachea (distal-proximal) axis to identify whether basal-like cells exhibited regional differences (Fig. 2a, b; Extended Data Fig. 3a, e). Tracheal basal cells at 12 weeks stain positively for TP63, KRT5, PDPN, EGFR, and F3 (Fig. 2b, Extended Data Fig. 3a, e), similar to tracheal basal cells in older lungs at 17 weeks of gestation (Fig. 1j), although basal cells in 12 week lungs are IL33 negative (Extended Data Fig. 3c). In the small cartilaginous airways, the majority of TP63+ cells were KRT5-, while KRT5+/TP63+ cells appeared to be present in raised patches of cells (Fig. 2b). In more distal regions of the airway, TP63 is detected and is co-localized with EGFR, while KRT5 is not detectable, F3 levels decline, and PDPN expression becomes more broad (Fig. 2c, Extended Data Fig. 3e). Protein localization of TP63 and KRT5 was confirmed by FISH for *TP63* and *KRT5* in 12 week fetal lungs, and mirrored patterns of protein expression along the proximal-distal axis (Extended Data Fig. 3a, b). By 17 weeks gestation, all TP63+ basal cells throughout the trachea and airway exhibit a mature “late” basal cell profile and expressed both TP63 and KRT5 (Extended Data Fig. 3d).

**Extended Data Figure 3.**
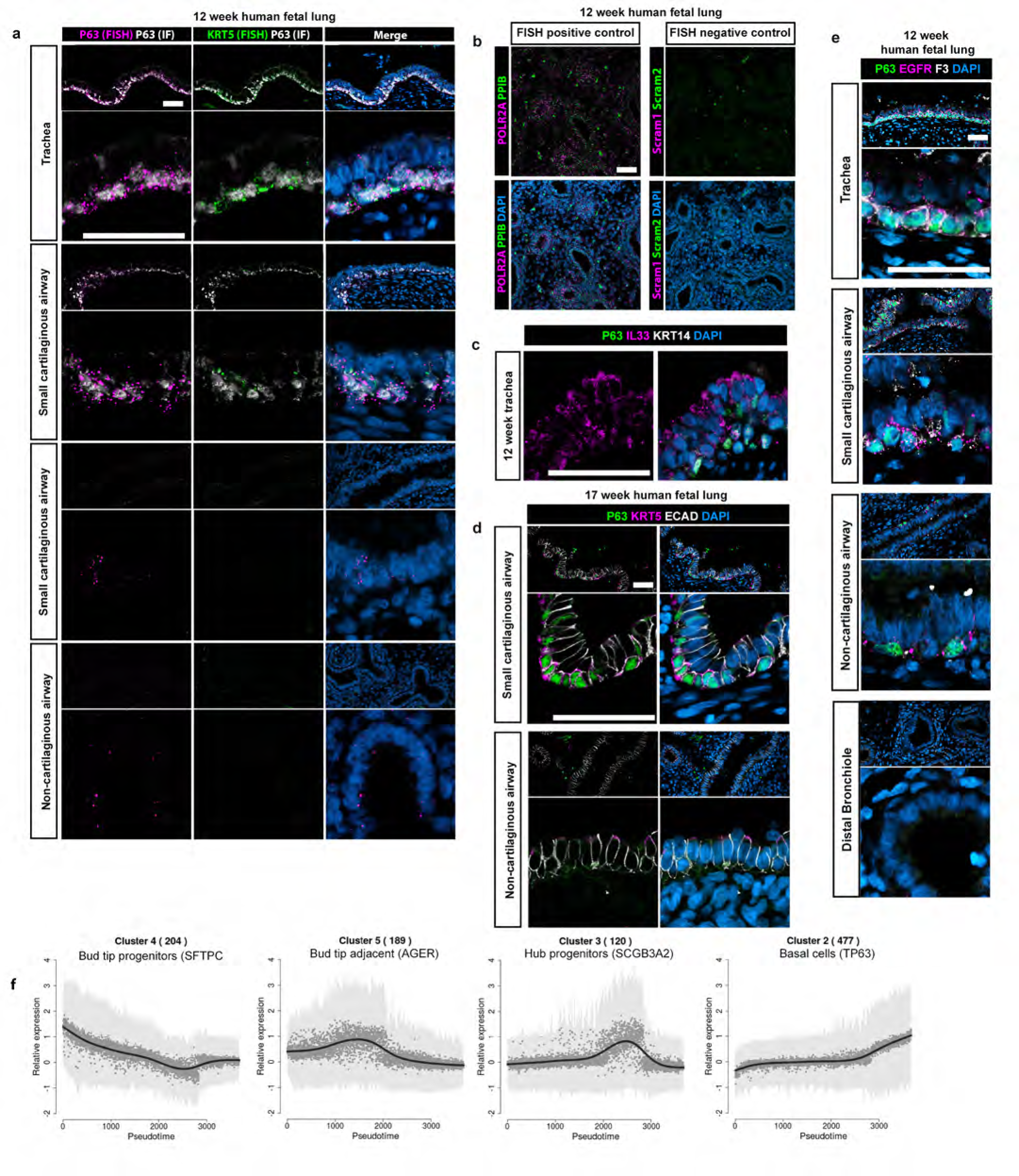
Fetal characterization by protein staining and mRNA *in situ* hybridization. **a**) mRNA *in situ* hybridization for TP63 (pink) and *KRT5* (green), combined with protein staining for TP63 (white) confirms the patterns seen in protein staining across the proximal-distal axis of the lung. Tracheal TP63+ basal cells exhibit strong *TP63* and *KRT5* mRNA expression. *KRT5* mRNA expression drops dramatically in the small cartilaginous airways and is absent in all *TP63*+ cells in the lower airways. Some cells exhibit low levels of *TP63* mRNA expression in the distal lung, including in the bud tip region. **b**) Positive (POLR2A [pink] and PPIB [green]) and negative control (Scrambled probes 1 [pink] and 2 [green]) stains for *in situ* hybridization. **c**) TP63+ (green) cells in the 12 week fetal trachea (n=3 biological replicates) do not stain positively for IL33 (pink) or KRT14 (white). **d**) By 17 weeks gestation (n=3 biological replicates), TP63+ cells are restricted to the trachea and large cartilaginous airways, and all TP63+ (green) cells exhibit KRT5 (pink) staining. **e**) To evaluate the phenotypic differences in protein expression of basal cell markers identified in Figure 1 across the proximal-distal axis in the developing 12 week fetal lung we stained for TP63 (green), EGFR (pink) and F3 (white). Similar to other markers, we observed strong co-staining for all 3 markers in tracheal basal cells, with a decrease in staining intensity for F3 to progressively more distal airways. Interestingly, EGFR appeared to mark TP63+ cells even in the non-cartilaginous airways, suggesting EGFR is an early marker for basal cells. Scale bars represent 50 µm. **f**) Average expression patterns of each variable gene cluster along pseudotime.

We next carried out analysis using SPRING to visualize continuous gene expression topologies, allowing us to infer lineage relationships using all combined fetal epithelial cells (Fig. 2c, d)^27^. This analysis inferred possible differentiation trajectories between cell populations, beginning with bud tip progenitors and transitioning through two major intermediate cell states, bud tip adjacent and hub progenitors, *en route* to the *TP63*+ basal cells (Fig. 2c). Cells from distal lung samples populated the bud tip progenitor and bud tip adjacent populations, whereas cells from the airways and tracheal scrapings populated several cell clusters, including hub progenitors, basal cells and differentiated airway cell types (Fig. 2c, d). To further support differentiation inferences from SPRING analysis, we used highly variable genes identified in bud tip progenitors, bud tip adjacent, hub progenitors and basal cells to order cells in pseudotime, which suggested a trajectory similar to that identified with SPRING (Fig. 2e, f). Average expression of variable genes were plotted to visualize fluctuations in gene expression across pseudotime and we determined that these variable genes could be classified into 5 major patterns (Extended Data Fig. 3f, Extended Data Table 2).

Hub progenitor cells, named for their central location within the SPRING plot, co-expressed high levels of *SCGB3A2*, *SFTPB* and *CFTR*(Fig. 2d). Expression patterns and anatomical location of hub progenitors were confirmed by co-expression of SCGB3A2 and SFTPB protein staining (Fig. 2b, Extended Data Fig. 4a, b). Anatomically, SCGB3A2+/SFTPB+ cells were most abundant in the non-cartilaginous and small to large cartilaginous airways at 12 weeks gestation and were physically located at the apical epithelium, close to the airway lumen. Hub cells were rare in the trachea (Fig. 2b, Extended Data Fig. 4a). By 18 weeks gestation, this population of cells was nearly undetectable by protein staining throughout the airway (Extended Data Fig. 4b) suggesting this population exists transiently. A population of uncommitted *Scgb3a2+/Upk3a*+ cells has been previously reported in the developing mouse airway, giving rise to club and multiciliated cells in the adult^28^. It is unclear if this murine population is equivalent to hub cells, which have not been previously reported in human lungs, but collectively, these data led us to hypothesis that hub progenitors may be a transitional progenitor population during human lung development.

**Extended Data Figure 4.**
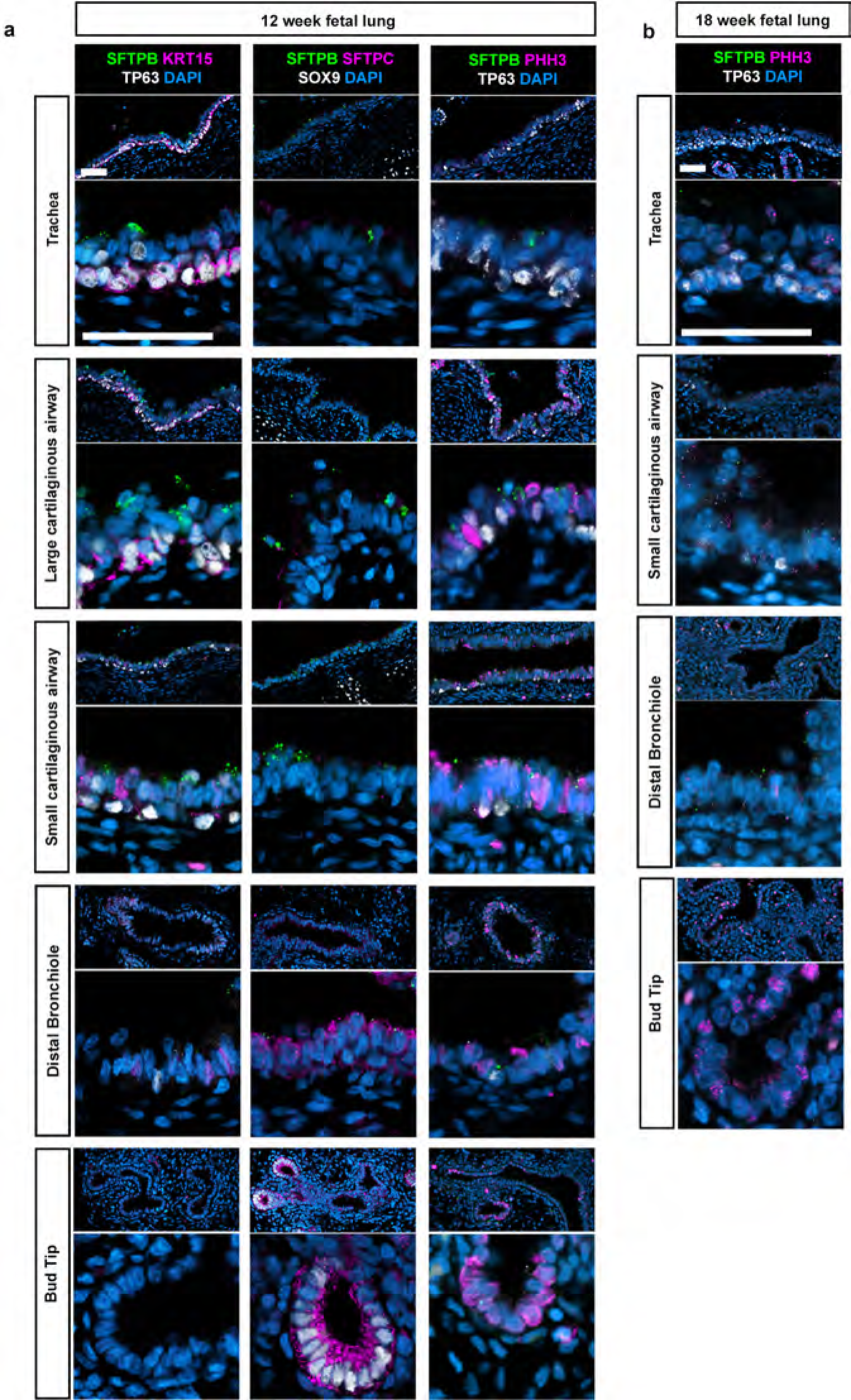
Characterization of hub progenitor cells. **a**) Protein staining characterization of hub progenitors throughout the proximal-distal axis of the 12 week fetal lung (n=3 biological replicates) shows that SFTPB+ (green) cells, which strongly mark hub progenitors by scRNAseq, are most prominent in the small and large cartilaginous airways at 12 weeks gestation. They are often apically localized, with some cells appearing above the epithelium (large cartilaginous airway left and second to left panels). Some hub progenitor cells appeared to contain weak nuclear TP63 (white) staining (left panel, small cartilaginous airway). To determine whether this apical localization was related to proliferation, we co-stained for SFTPB (green) and Phosphohistone H3 (pink), third panel from left. This staining showed that some, but not the majority, of hub progenitors were proliferative. Scale bar represents 50 µm. **b**) By 18 weeks gestation (n=3 biological replicates), hub progenitors (SFTPB, green) were largely absent, suggesting this population is transitory during development. The lung epithelium at 18 weeks was also substantially less proliferative than at 12 weeks as determined by phospho-histone H3 staining (PHH3, pink). Scale bars represent 50 µm.

In order to understand possible mechanisms by which bud tip progenitors are instructed to differentiate into TP63+ basal stem cells, we performed an analysis of cell signaling and transcription factor networks, and found that TGFβ signaling pathway genes were significantly enriched in the pseudotemporally variable genes (Fisher’s exact test, nominal P=0.00082; Fig. 2g). Interestingly, inhibition of SMAD-mediated TGFβ and BMP signaling has been implicated in regulating basal cell differentiation in adult lung regeneration^4,21,22,26,29^, whereas our analysis showed that TGFβ and BMP receptor expression is enriched in the bud tip and bud tip adjacent clusters, but expression is reduced in hub progenitor and basal cells (Fig. 2g). We therefore hypothesized that transient activation of SMAD signaling may induce airway lineages including basal stem cells from fetal bud tip progenitors. Consistent with this hypothesis, protein staining in fetal lung tissue sections revealed weak nuclear phospho-SMAD staining in bud tip progenitors (Extended Data Fig. 5a) whereas immature TP63+ cells in non-cartilaginous and small-cartilaginous airways exhibited strong nuclear staining of phospho-SMAD2 (pSMAD2; Fig. 2h) and phospho-SMAD1,5,8 (pSMAD1,5,8; Extended Data Fig. 5b) at 12 weeks gestation, suggesting active SMAD signaling in these cells. Consistent with previous data in mature basal cells within the adult human trachea^1,2,26^, mature TP63+ basal cells in the 12 week trachea exhibited very weak nuclear pSMAD2 and pSMAD1,5,8 staining, but adjacent luminal tracheal epithelial cells had strong nuclear pSMAD expression (Fig. 2h, Extended Data Fig. 5b). Expression of the co-SMAD, SMAD4, was similar throughout the lung epithelium at all time points (Extended Data Fig. a, b, c). Nuclear pSMADs are associated with basal cells only at earlier developmental stages (i.e. 12 weeks gestation), as TP63+ basal cells observed in 17 week fetal lungs lacked strong nuclear pSMAD staining irrespective of anatomical location (Extended Data Fig. 5c). These *in vivo* observations supported the hypothesis that transient activation of SMADs may induce a basal cell state, whereas active SMAD signaling is excluded from mature basal cells.

**Extended Data Figure 5.**
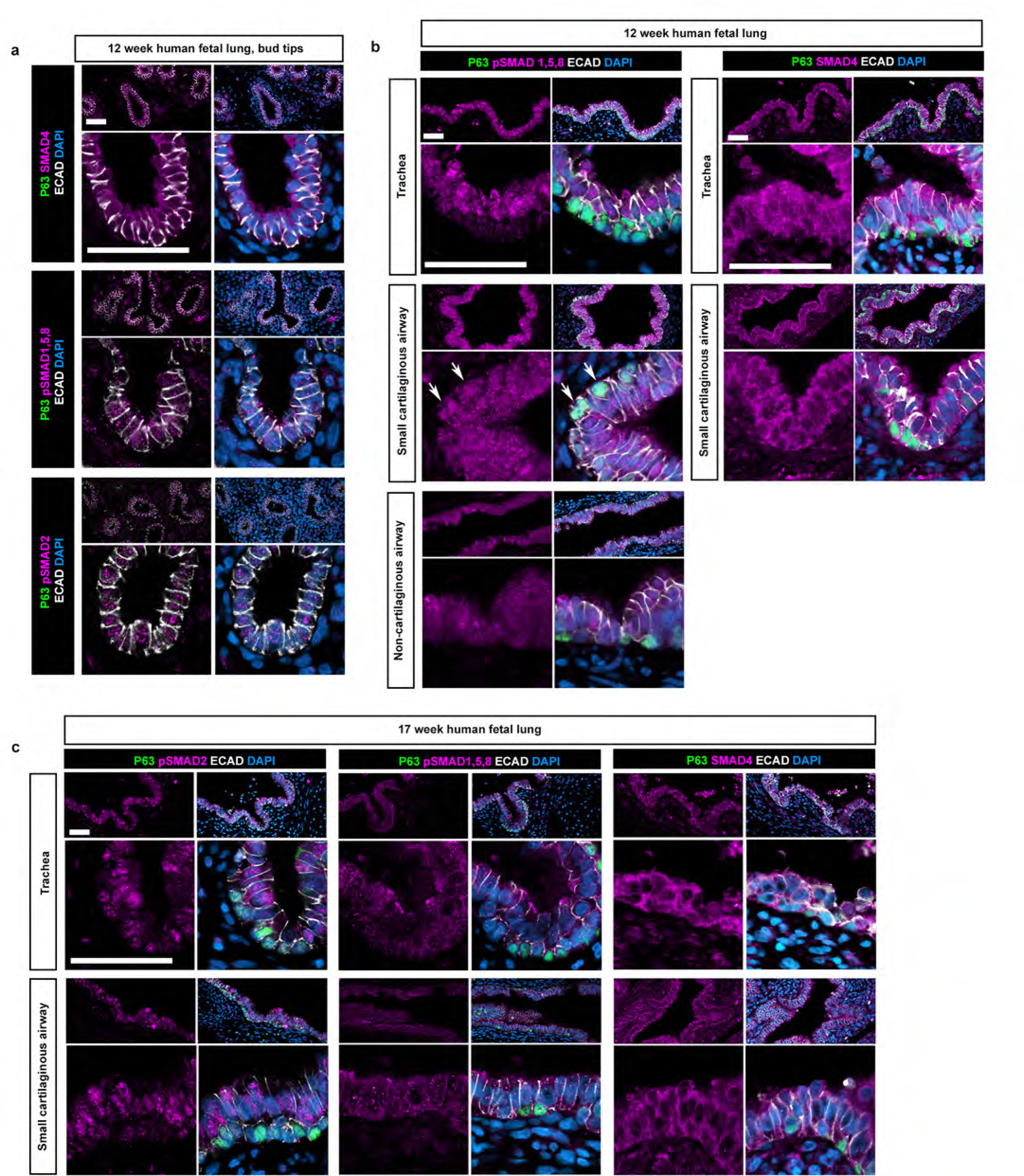
Characterization of SMAD signaling in the developing fetal lung. **a**) staining for pSMAD4 (pink; top panels), pSMAD1,5,8 (pink; middle panels) and SMAD4 (pink; bottom panels) with TP63 (green) shows that nuclear SMAD staining is low in the bud tip progenitors. ECAD staining marks the cell membranes (white). Scale bars represent 50 µm. **b**) Staining for pSMAD1,5,8 (pink) and TP63 (green) across the proximal-distal axis of the developing lung shows very weak nuclear pSMAD1,5,8 staining in the non-cartilaginous airways, but stronger nuclear staining in TP63+ cells within the small cartilaginous airways. In the trachea, TP63+ cells contain very little to no nuclear pSMAD1,5,8. This pattern is similar to the pattern seen for pSMAD2 (Fig. 2h). Staining for SMAD4, a non-phosphorylated binding partner for both SMAD2 and SMAD1,5,8, is present in all epithelial cells. ECAD staining marks the cell membranes (white). Scale bars represent 50 µm. **c**) Staining for pSMAD2 (pink; left panels), pSMAD1,5,8 (pink; middle panels) and SMAD4 (pink; right panels) with TP63 (green) show that, by 17 weeks gestation, all TP63+ cells exhibit little to no nuclear pSMAD staining. ECAD staining marks the cell membranes (white). Scale bars represent 50 µm.

### TGFβ1 and BMP4-mediated SMAD activation induces functional basal stem cells from bud tip progenitors in vitro

To test the hypothesis that transient SMAD activation promotes differentiation of bud tip progenitors towards a basal cell lineage in human tissue, we used human fetal bud tip progenitor organoids isolated from 12 week fetal lungs^11,30^ that were maintained as progenitors in culture, and tested if activators or inhibitors of the TGFβ/BMP signaling pathways alone, or in combination, influenced *TP63* expression (Fig. 3a, Extended Data Fig. 6a-c).

**Figure 3.**
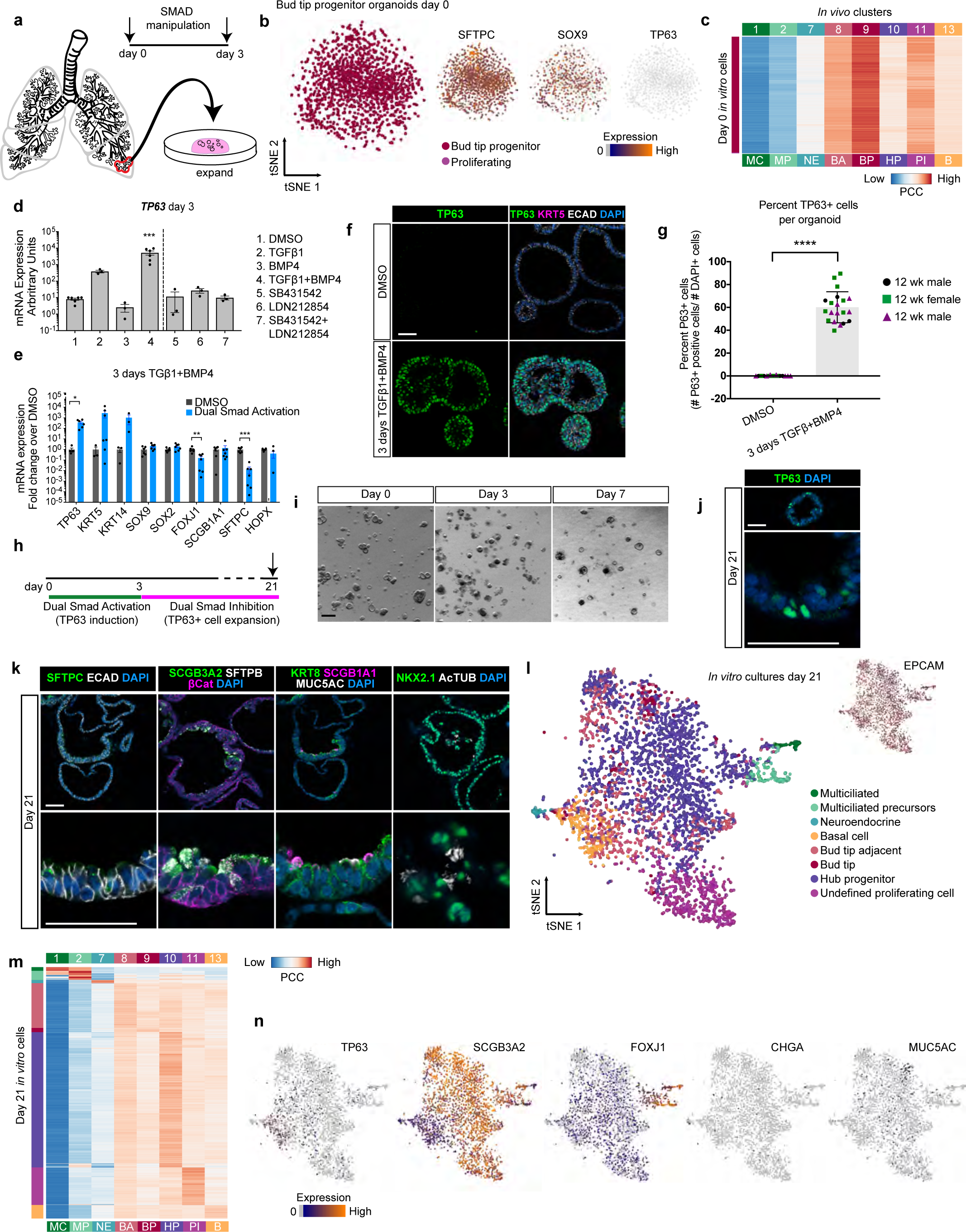
SMAD activation induces a bud tip to airway/basal cell transition in bud tip progenitor organoids. **a**) Schematic of experimental design. All experiments were carried out using n=3 12 week human fetal lung-derived bud tip progenitor organoids. **b**) Bud tip progenitor organoids maintained in progenitor maintenance medium were collected for scRNAseq prior to treatment with DSA (day 0). tSNE clustering (left) and feature plots (middle, right) of *SFTPC*, *SOX9* and *TP63.***c**) The expression profile of each day 0 *in vitro* cell was compared to expression profiles of all *in vivo* epithelial cell clusters and assigned a correlation score. The majority of cells at day 0 were highly correlated with *in vivo* bud tip progenitor cells or proliferating cells. **d**) Organoids were treated for 3 days with SMAD activation or inhibition conditions and expression of *TP63* was evaluated by QRT-PCR for all treatment groups. Treatment for 3 days with both TGF 1 (100 ng/mL) and BMP4 (100 ng/mL) led to a significant increase in *TP63* expression relative to all other groups (One-way ANOVA alpha=0.05, F=21.19, R square=0.9008, p<0.0001; Tukey’s multiple comparisons of the mean of each group versus the mean in all other groups, p values are reported on the graph. p<0.05= * ; p<0.01 = **, p<0.001 = ***, p<0.0001 = **** ; 3 days TGF01 and BMP4 referred to as ‘dual SMAD activation’, or ‘DSA’). Data is plotted as arbitrary units. Error bars are plotted to show mean +/− the standard error of the mean. N=3 independent biological specimens. Data is from a single experiment and is representative of n=3 experiments. **e**) QRT-PCR for markers of canonical differentiated lung epithelial cell types showing DMSO (gray bars) and DSA treated (blue bars) organoids after 3 total days of treatment. Data is plotted as fold change over DMSO controls. DSA treated organoids exhibited a 370-fold increase over DMSO controls in mean *TP63* expression, a 2780-fold increase in mean KRT5 expression and a 1045-fold increase in mean *KRT14* expression, all basal canonical cell markers. *TP63* expression was statistically significantly higher than DMSO controls (Two-sided Mann-Whitney Test, p=0.0238), but the increases in KRT5 and KRT14 were not statistically significant. No other markers exhibited increases in expression after 3 days of DSA treatment. Some markers exhibited a significant reduction in expression, including the ciliated cell marker FOXJ1 (Two-sided Mann-Whitney Test, p=0.02) and bud tip progenitor marker/AECII marker SFTPC (Two-sided Mann-Whitney Test, p=0.002). Error bars represent the mean +/− the standard error of the mean. n=3 independent biological specimens, and data is from a single experiment and is representative of n=3 experiments. **f**) Protein staining of DMSO treated (control) fetal bud tip progenitor organoids (top row) and 3 days of DSA treatment (bottom row) for TP63+ staining (green), KRT5 (pink) and DAPI (blue). Scale bar represents 50 µm. **g**) Quantification of (f). Total number of TP63+ cells were counted for 3-9 individual organoids across 3 biological replicates. DMSO treated controls exhibited 0.125% (+/− 0.08%) TP63+ cells, whereas 60.13% (+/− 3.035%) of cells within organoids treated with 3 days of TGF/1 and BMP4 showed positive TP63 staining. n=3 independent biological specimens. **h**) Experimental schematic for panels i-n. **i**) Dual SMAD inhibition (DSI) led to organoid expansion. **j**) After 21 days, organoids contained TP63+ (green) cells. Scale bars represent 50 µm). **k**) After 21 days in culture, organoids were interrogated for protein staining of the bud tip progenitor marker SFTPC (green), hub progenitor cells (SCGB3A2 [green], SFTPB [white]), club cells (SCGB1A1 [pink]), goblet cells (MUC5AC [white]) and multiciliated cells (AcTUB [white]). **l**) 21 day organoids were subjected to scRNAseq and visualized by tSNE. Cell identities were based on cell IDs as described in Fig. 1. **m**) The expression profile of each *in vitro*-derived cell was compared to expression profiles of all *in vivo* epithelial cell clusters and assigned an identity based on correlation tp an *in vivo* cluster. **n**) Feature plots of individual genes for markers of cilitated cells (*FOXJ1*), neuroendocrine cells (*CHGA*), basal cells (*TP63*), and hub progenitor cells (*SCGB3A2*).

**Extended Data Figure 6.**
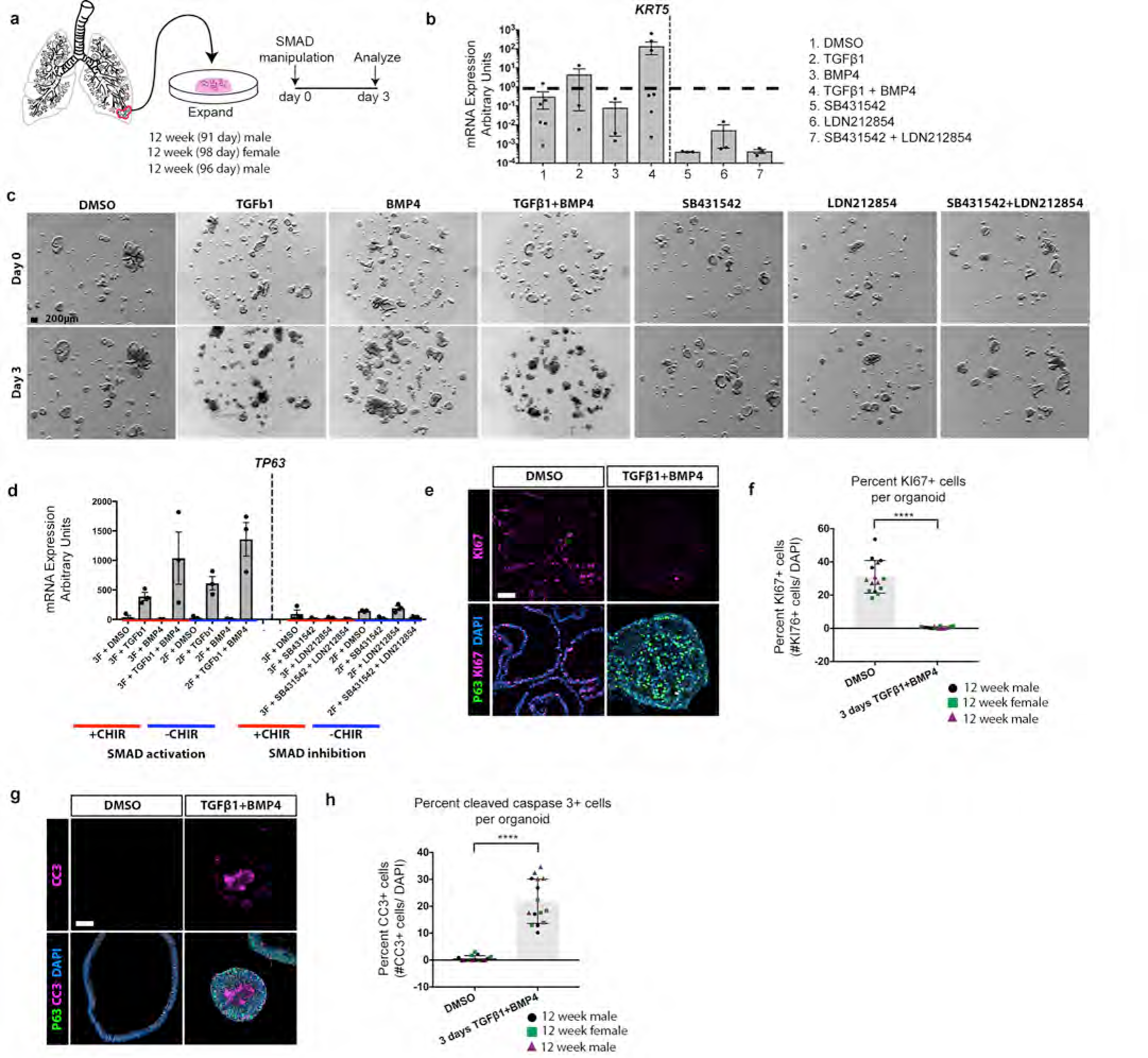
Screen for growth factor combinations that induce *TP63* expression in fetal bud tip progenitor organoids. **a**) Schematic of experimental design. Bud tip progenitors were enzymatically and mechanically isolated from n=3 independed 12 week human fetal lung specimens for all experiments as previously described^11^ and expanded as 3-dimensional organoids in Matrigel droplets. Once cultures were established, they were treated with serum-free progenitor maintenance medium supplemented with combinations of activators and/or inhibitors of TGF/ and BMP signaling. Organoids were collected after 3 days in culture and analyzed for mRNA and protein expression. **b**) QRT-PCR for *KRT5* showed no significant changes in expression for any group (one-way ANOVA, p>0.05). Data is presented as mean +/− the standard error of the mean. **c**) Brightfield images of a single well of organoids for each group at day 0 and after 3 days of treatment. Treatment with TGFi1 alone or TGF1 and BMP4 led to a reduction in organoid growth, an increase in apparent density of the organoids, and the appearance of many dead cells and debris around the organoids. **d**) We tested whether the presence of CHIR99021, a ROCK inhibitor and potent activator of the WNT pathway, affected the outcome of SMAD manipulation on *TP63* expression. No statistical differences were observed between groups treated in the presence or absence of CHIR99021, though we note that variability within groups is high (one-way ANOVA, p>0.05). For this work, we continued to include CHIR99021 in the dual SMAD activation (DSA) medium because it seemed to improve survival of organoid cultures. Data is reported as arbitrary units and error bars represent the mean +/− the standard error of the mean. **e**) Protein staining of proliferation marker KI67 (pink) and TP63 (green) in organoids treated with DMSO or with 3 days of DSA shows that DMSO treated organoids are very proliferative and do not contain any TP63+ cells, whereas organoids treated with 3 days of DSA medium exhibit very few proliferating cells and many TP63+ cells. DSA treated organoids also exhibit a denser morphology. Scale bars represent 50 µm. **f**) Quantification of e. Total number of KI67+ cells were counted for 4-5 individual organoids across 3 biological replicates. DMSO treated controls exhibited 31.04% (+/− 9.79%) KI67+ cells, whereas 0.54% (+/− 2.53%) of cells within organoids treated with 3 days of DSA showed positive KI67 staining. DSA treated organoids exhibited a statistically significant decrease in the number of KI67+ cells (Two-sided Mann-Whitney test, P<0.0001). **g**) Protein staining of apoptosis marker Cleaved Caspase 3 (CC3; pink) and TP63 (green) in organoids treated with DMSO or with 3 days of DSA shows that DMSO treated organoids do not exhibit any apoptosis staining and do not contain any TP63+ cells, whereas organoids treated with 3 days of DSA medium exhibit a dramatic increase in CC3+ cells and many TP63+ cells. DSA treated organoids also exhibit a denser morphology. Scale bar represent 50 µm. **h**) Quantification of g. Total number of CC3+ cells were counted for 5 individual organoids across 3 biological replicates. DMSO treated controls exhibited 0.64% (+/− 0.25%) CC3+ cells, whereas 21.86% (+/−2.12%) of cells within organoids treated with 3 days of DSA showed positive CC3 staining. DSA treated organoids exhibited a statistically significant increase in the number of CC3+ cells (Two-sided Mann-Whitney test, p<0.0001).

First, in order to define cell composition in cultured bud tip organoids, we analyzed the transcriptomes of 1,592 cells prior to differentiation experiments (‘day 0’) and compared these results to the scRNAseq data from *in vivo* fetal lung epithelial cells. Cell clustering of the ‘day 0’ bud tip progenitor organoids using tSNE revealed 2 populations of cells; bud tip progenitor cells and proliferating cells (Fig. 3b). Each cell from the *in vitro* day 0 bud tip organoid sample was compared to the *in vivo* epithelial cell gene expression signatures (as identified in Fig. 1), and was assigned a correlation score based on similarity to *in vivo* cell clusters (Fig. 3c). *In vitro* maintained bud tip progenitor cells shared the highest degree of gene expression similarity with *in vivo* bud tip progenitor cells and proliferating cells (Fig. 3c). Plots for individual genes reveal the majority of cells at day 0 are *SFTPC*+ and *SOX9*+ but negative for markers of more differentiated cell types, including the basal cell marker *TP63* (Fig. 3b).

Supplementing normal progenitor organoid growth medium with SMAD activators or repressors for 3 days revealed that TGFβ1 plus BMP4 (herein referred to as ‘dual SMAD activation’; DSA) led to the most significant increase in *TP63* expression by QRT-PCR (Fig. 3d). 3 days of DSA also led to non-significant increases of mRNA expression over untreated controls in canonical basal cell markers *KRT5* and *KRT14*, but showed no increase in markers for other lung epithelial cell types (Fig. 3e). DSA significantly increased *TP63* expression both in the presence and absence of CHIR99021 (Extended Data Fig. 6d), a GSK3β inhibitor that is required to maintain bud tip organoids in their undifferentiated state^5-8,11^. Protein staining further revealed that 60.13% (+/−13.04%) of DSA treated bud tip organoid cells expressed TP63 after 3 days, compared to 0% in controls (Fig. 3f, g). DSA treatment led to more dense epithelial structures (Fig. 3f, i, Extended Data Fig. 6c, e, g) with significantly reduced proliferation as measured by KI67 staining (Extended Data Fig. 6e, f) and increased apoptosis (Extended Data Fig. 6g, h). Consistent with bud tip progenitor organoid experiments, we similarly found that when explanted pieces of whole distal lung tissue (10-11 week fetal lung) were treated with DSA, significant increases in TP63 protein and mRNA expression were observed (Extended Data Fig. 7a-d), suggesting that DSA is a potent inducer of TP63 in multiple contexts.

**Extended Data Figure 7.**
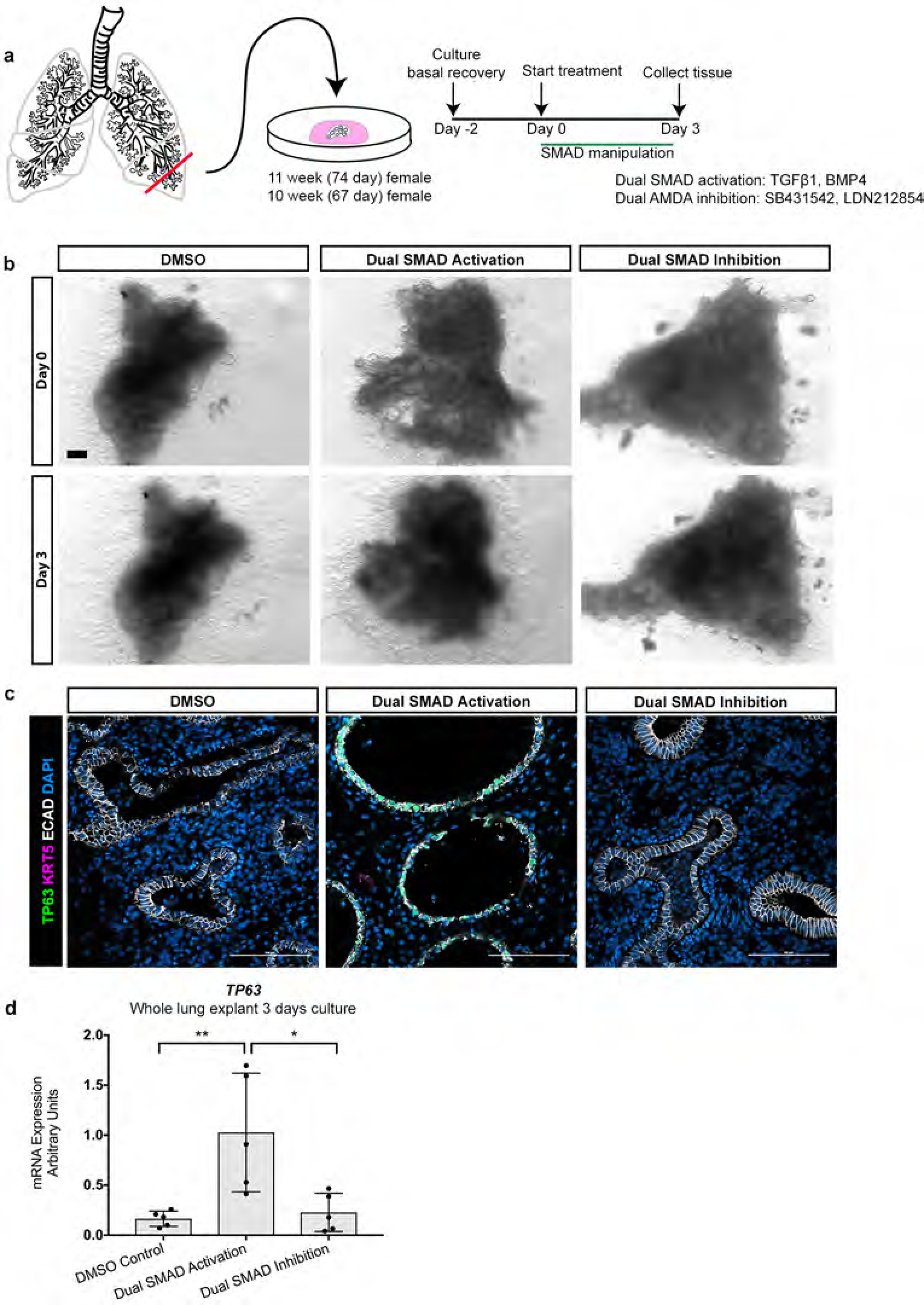
Fetal lung explant cultures. **a**) Schematic of experimental design. The most distal portions of a 10 week and 11 week fetal lung were cut with a scalpel to 1cm^2^ units and laid on top of a fresh Matrigel droplet prior to solidification (see methods). They were cultured in serum-free, growth factor-free medium supplemented with 1% Fetal Bovine Serum for 2 days to facilitate recovery. They were then treated with serum-free medium (no addition of bud tip progenitor maintenance growth factors) supplemented with TGF and BMP pathway activators and inhibitors. **b**) Bright field images of lung explants on day 0 (2 days post plating) and day 3 of treatment. All explants survived and expanded in culture. DSA appeared to reduce the amount of epithelium and allow expansion of the mesenchyme, whereas DSI appeared to allow expansion of the epithelium and the mesenchyme. Scale bar represents 1 mm. **c**) Protein staining for TP63 (green) and KRT5 (pink) reveals a drastic increase in the number of TP63+ cells in the DSA tissue compared to the DMSO and DSI groups. DSA treatment also led to a reduction in the total amount of epithelium and a morphologically more thin epithelium compared to other groups. KRT5 was not detected in any group. Scale bars represent 100 µm. **d**) 2 explants from the 10 week lung (technical replicates) and 3 explants from the 11 week lung (technical replicates) were processed for RNA extraction. QRT-PCR of *TP63* showed that 3 days of DSA treatment led to a significant increase in *TP63* expression relative to both DMSO controls and to DSI treated organoids (one-way ANOVA; Multiple comparisons comparing the mean of each group to the mean of every other group, p<0.001 between DMSO controls and DSA, p<0.05 between DSA and DSI treated organoids). There was no statistical difference in *TP63* expression between DMSO controls and DSI treated organoids. Error bars represent the mean +/− the standard error of the mean.

Our findings show that TP63 is robustly induced in bud tip progenitor cells exposed to DSA but that prolonged treatment also blocked proliferation and induced cell death, creating technical challenges to isolating and studying this population. Previous work in mice and humans has shown that inhibition of TGFβ/BMP (dual SMAD inhibition; DSI) is required for expansion of mature adult basal stem cells in culture^21,26,29^. Based on our data and published work, we reasoned that, while DSA is sufficient to induce TP63 expression, it may be detrimental to long-term growth and cell expansion. Therefore, we screened for growth factor conditions that allowed the DSA-induced cell population to expand in culture (Fig. 3h-i; Extended Data Figure 8a-e). We found that supplementation of the medium with FGF10 and Y27632, a RHO kinase inhibitor, in addition to inhibitors of TGFβ and BMP (‘DSI expansion medium’: FGF10, A8308, NOGGIN, Y27632), promoted organoid survival and expansion, and was permissive for continued TP63 expression (Fig. 3i, j; Extended Data Fig. 8a-e). DSI expansion medium is consistent with published reports demonstrating the importance of these factors for basal cell proliferation, survival and/or expansion^9,10,26,29,31^. Organoids grown in expansion medium also demonstrated increased mRNA expression of markers canonically associated with multiciliated and club cell lineages (*FOXJ1, SCGB1A1,* respectively (Extended Data Fig. 8e) and protein expression for markers of hub progenitors (SCGB3A2, SFTPB) multiciliated (FOXJ1, ACTUB), club (SCGB1A1), goblet (MUC5AC) and neuroendocrine (CHGA, SYN; Fig. 3k, Extended Data Fig. 9b). After several days in culture, many beating multiciliated cells were present within organoids and the proteinaceous luminal contents within organoids appeared to swirl with directionality, suggesting that multiciliated cells are functional and able to propel luminal contents (Extended Data Video 1).

**Extended Data Figure 8.**
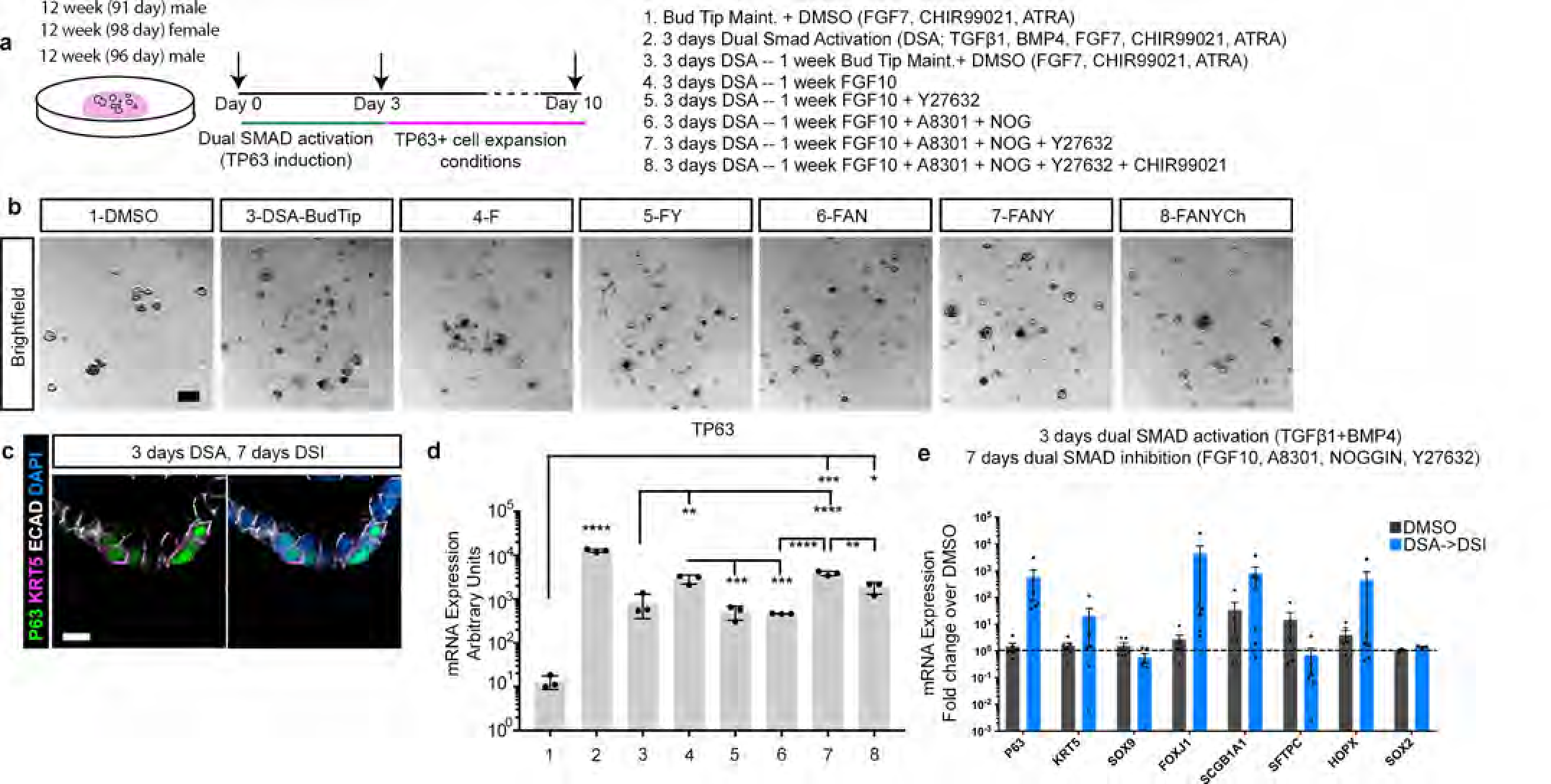
Screen for factors that maintain growth and expansion of TP63+ cells in culture. **a**) Experimental schematic. Fetal bud tip progenitor organoids were treated with DSA for 3 days and then treated with various combinations of growth factors for an additional 7 days to identify conditions that maintained and expanded TP63+ population of cells (10 days total). Experimental conditions are listed as groups 1-8. **b**) Brightfield images of organoids 10 days after treatment. Treatment with group 7 medium led to the best expansion and survival of organoids. Scale bar represents 500 µm. **c**) After 7 days of expansion in group 7 medium (Dual SMAD Inhibition; DSI), organoids maintained cells that expressed TP63 (green), and a very few number of cells exhibited staining for mature basal cell marker KRT5 (pink). ECAD marks the cell membrane (white). No cells expressed other basal cell marker PDPN (pink), nor did any cells express the bud tip adjacent/AECI marker RAGE (white). Scale bar represents 50 µm. **c**) Expression of *TP63* was evaluated by QRT-PCR for all treatment groups. Treatment for 3 with DSA led to a statistically significant increase in TP63 expression over DMSO controls. Treatment with DSA followed by 7 days of treatment with medium in group 7 (FGF10, A8301, NOGGIN, Y2763) or group 8 (FGF10, A8301, NOGGIN, Y2763, CHIR99021) medium led to a significant increase in *TP63* expression compared with the DMSO control (One-way ANOVA, alpha=0.05, F=191.9, R square=0.9882, p<0.0001; Tukey’s multiple comparisons of the mean of each group versus the mean in all other groups, p values are reported on the graph. p<0.05= * ; p<0.01 = **, p<0.001= ***, p<0.0001= ****). Data is plotted as arbitrary units. Error bars are plotted to show mean +/− the standard error of the mean. Data is from a single experiment and is representative of n=3 experiments. **e**) QRT-PCR for markers of canonical differentiated lung epithelial cell types showing DMSO (gray bars) and DSA––DSI treated (blue bars) organoids after 3 days DSI and 7 days DSI treatment. Data is plotted as fold change over DMSO controls. DSA––DSI treated organoids exhibited a 394-fold increase over DMSO controls in mean *TP63* expression. No markers exhibited statistically significant increases or decreases in expression after treatment, although trends suggest an increase in *TP63,* ciliated cell marker *FOXJ1* and club cell marker *SCGB1A1*. Error bars represent the mean +/− the standard error of the mean.

To quantify and evaluate the transcriptomic similarity between *in vitro* derived cells and *in vivo* fetal lung epithelial cells, we performed scRNAseq on 3,741 cells from bud tip progenitor organoids treated for 3 days with DSA followed by 18 days of DSI expansion medium. tSNE visualization of clusters from day 21 organoids revealed 8 different airway cells clusters (Fig. 3l), which showed a high degree of correlation with *in vivo* cell signatures for hub progenitors, bud tip progenitors, bud tip adjacent, basal cell, neuroendocrine, proliferating and multiciliated cell signatures (Fig. 3m). Cluster identities were further supported by the expression of marker genes for each population (Fig. 3n). Interestingly, while certain cells expressed markers of mature goblet (MUC5AC) and club (SCGB1A1) cells (Fig. 3n), which were also observed via protein staining (Fig. 3k, Extended Data Fig. 9b), the transcriptomic profile of these cells did not fully match to *in vivo* signatures of human fetal goblet and club cells (Fig. 3m). Instead, these secretory cells matched closest to hub progenitors, suggesting they remain immature at this time point. *In vitro*-derived *TP63+* cells also express *KRT15*, but low levels of *KRT5*, suggesting these cells have an “early” basal cell-like phenotype.

**Extended Data Figure 9.**
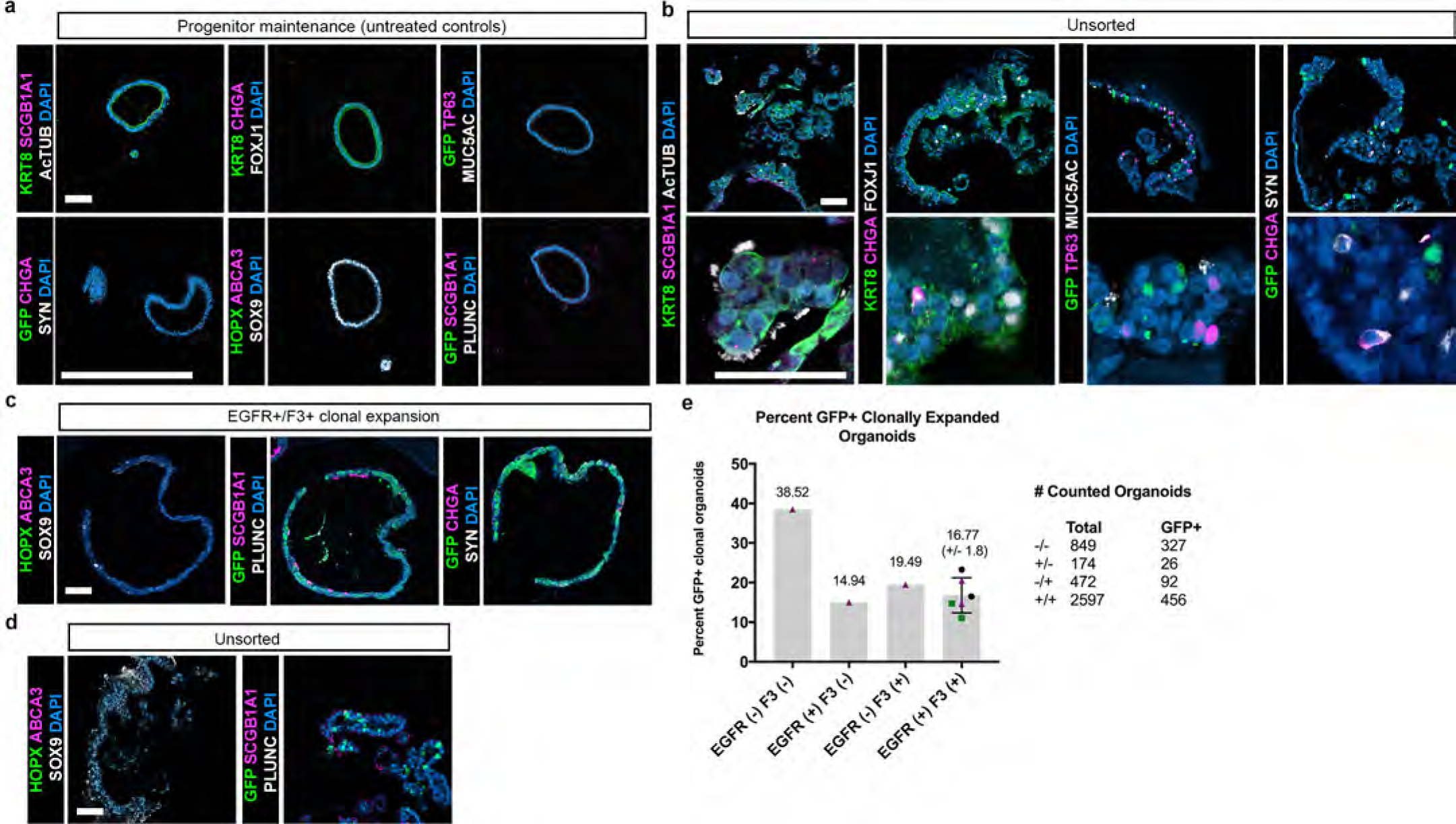
Sorting and clonal expansion of fetal bud tip progenitor organoids. **a**) Bud tip progenitor organoids from n=3 biological replicates from 12 week fetal lungs were maintained in serum-free progenitor maintenance medium (FGF7, CHIR99021, ATRA) for 56 days. Staining for markers of differentiated lung epithelial cell types determined that bud tip progenitor organoids did not contain any differentiated cell types (club cell marker SCGB1A1 (pink), neuroendocrine markers Chromagranin A (CHGA; pink) and synaptophysin (SYN; white), ciliated cell marker FOXJ1 (white), basal cell marker TP63 (pink), goblet cell marker mucin 5AC (MUC5AC; white), AECII marker ABCA3 (pink), AECI and hub progenitor cell marker HOPX (green), secretory lineage marker PLUNC (white)). The majority of bud tip progenitor organoid cells were SOX9+ (white). Scale bar represents 100 µm. **b**) Organoids that had been infected with GFP lentivirus but not sorted and left to expand in basal cell expansion medium were collected after 56 days in culture and stained for differentiated epithelial cell markers. Unsorted organoids exhibited clear positive staining for club cell marker SCGB1A1 (pink) and multiciliated markers Acetylated Tubulin (white) and FOXJ1 (white). Neuroendocrine cells were clearly detected (CHGA, pink; SYN, white). Organoids also exhibited TP63+ cells (pink) and cells that stained positive for goblet cell marker MUC5AC (white). Scale bar represents 50 µm. **c**) Organoids treated for 3 days DSA and 18 days DSI-expansion, FACS sorted for EGFR/F3 and clonally expanded in DSI expansion medium were stained for differentiated cell markers. Staining for AECI and hub progenitor marker HOPX (green) was negative, as was staining for AECII marker ABCA3 (pink). Organoids grew clonally, with organoids being either entirely GFP negative or GFP positive (GFP, green, second panel). Many cells exhibited positive staining for club cell marker SCGB1A1 (pink), but staining for secretory lineage marker PLUNC (white) was undetected. In EGFR+/F3+ clonal organoids, no neuroendocrine cells were detected (CHGA, pink; SYN, white), though these cells were clearly detected in unsorted organoids. Scale bar represents 50 µm. **d**) Unsorted organoids also did not show any positive staining for AECI/hub progenitor marker HOPX (green), or AECII marker ABCA3 (pink). Consistent with results from sorted organoids, the secretory lineage marker PLUNC was not detected (white). Scale bar represents 50 µm. **e**) Graph of the percent of GFP+ organoids versus total number of organoids from each group. The total number of organoids counted per group is reported. 2 wells of multiple organoids were counted for each biological replicate for the EGFR+/F3+ group.

### *In vitro*-derived TP63+ cells are functional basal stem cells

Lastly, we asked whether *in vitro* derived TP63+ cells exhibited functional hallmarks of basal stem cells, including clonal expansion, self-renewal, and multilineage differentiation. In order to determine if DSA-induced TP63+ cells have the capability for self-renewal and multi-lineage differentiation, we isolated TP63+ cells with FACS using the basal cell enriched cell surface proteins EGFR and F3, which are expressed in TP63+ cells in native basal cells and in TP63+ cells from DSA-DSI expansion treated organoids (Fig. 1j, Fig. 4a, b, c Extended Data Fig. 10a). Prior to induction of TP63 by DSA, bud tip organoids were infected with a lentivirus to drive constitutive GFP expression in a random subset of cells, (∼20% efficiency, Extended Data Fig. 10c, d) in order to track and visualize these cells over time. Bud tip organoids were then treated with DSA for 3 days to induce *TP63* expression, and expanded for 18 days in DSI expansion medium (Fig. 4a). Organoids were subsequently dissociated and FACS was used to isolate EGFR+/F3+ cells (Fig. 4c, Extended Data Fig. 10e-h). FACS isolated cells were immediately affixed to glass slides via cytospin and were stained for TP63 protein. This analysis showed that 92.09 (+/− 1.66)% of EGFR+/F3+ cells co-expressed nuclear TP63, whereas EGFR+/F3−, EGFR−/F3+ or EGFR−/F3− fractions had a far lower proportion of cells that expressed TP63 (Fig. 4e). After sorting, EGFR+/F3+ cells were replated and grown *in vitro* in DSI expansion medium, where approximately 15-30% of all organoids across groups were GFP+ (Extended Data Fig. 9e). Whole organoids were composed either entirely of GFP+ or GFP− cells, but no organoids contained a mixture of GFP+ and GFP− cells, suggesting that individual organoids were derived from clonal expansion rather than from cell aggregation (Fig. 4f, Extended Data Fig. 9c, e). Clonally expanded EGFR+/F3+ cells gave rise toTP63+ basal, MUC5AC+ goblet-like, SCGB1A1+ club-like and AcTUB+/FOXJ1+ multiciliated cells as shown by protein staining (Fig. 4g, Extended Data Fig. 9c), which were not present in untreated controls (Extended Data Fig. 9a). Interestingly, no neuroendocrine cells were detected in EGFR+/F3+ clonally expanded basal cell-derived organoids, suggesting either that neuroendocrine cells are derived from an earlier progenitor or that there is heterogeneity within the basal cell population, with certain basal cells able to give rise to this restricted lineage. Further, clonally expanded GFP+ organoids exhibited functional multiciliated cells that beat, along with swirling luminal contents (Extended Data Video 2). Together, this data supports the hypothesis that DSA induces a population of functional basal cells from bud tip progenitor organoids *in vitro*.

**Figure 4.**
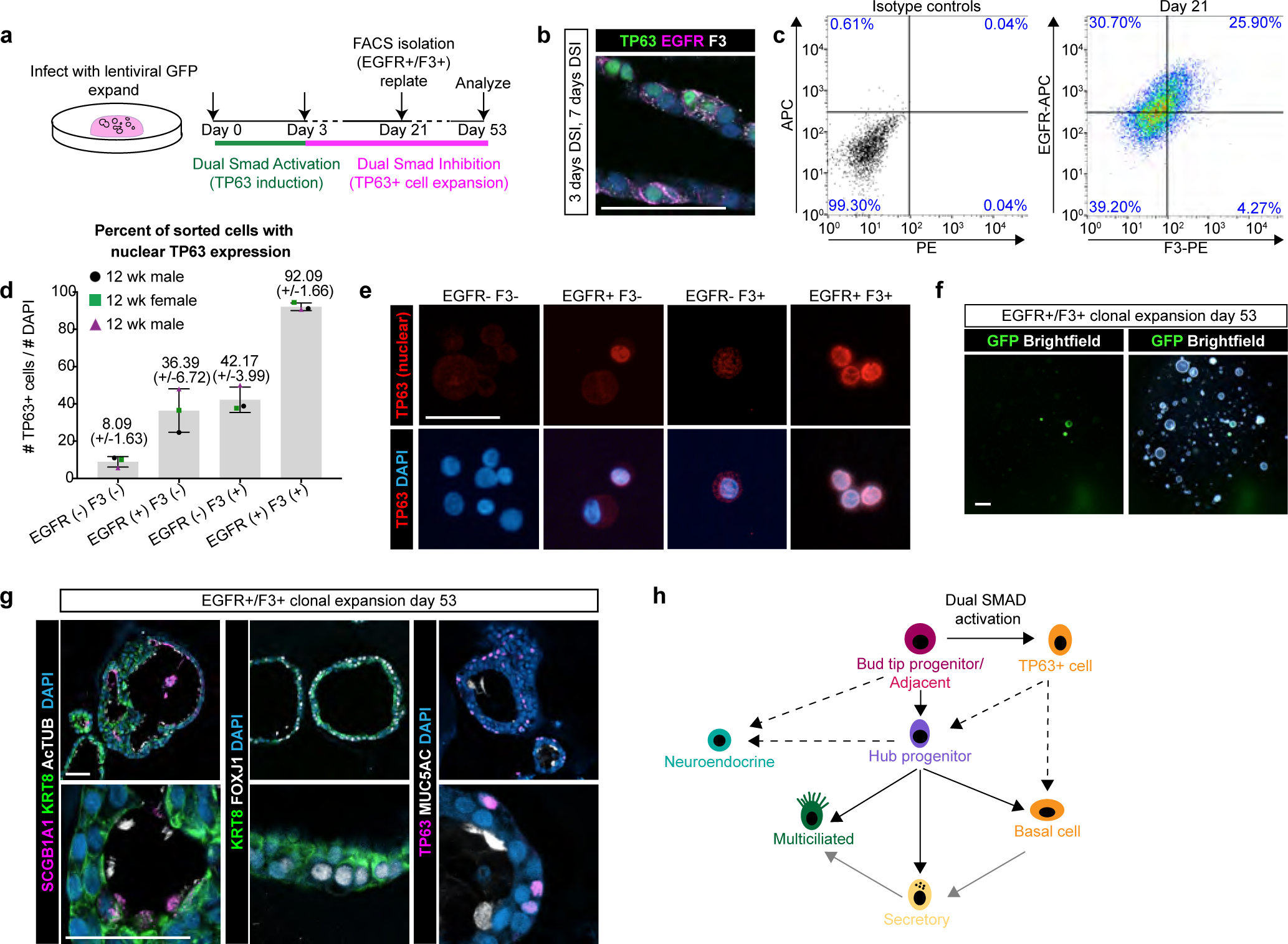
*In vitro*-derived TP63+ cells are functional basal stem cells. **a**) Experimental overview of cell sorting and clonal expansion experiments. 12 week fetal bud tip progenitor organoids (n=3 biological replicates) were infected with a lentiviral construct to express constitutive GFP. Organoids were then treated with 3 days DSA to induce *TP63* expression, followed by 18-19 days of treatment with DSI expansion medium, Fluoresence Activated Cell Sorting (FACS) to isolate TP63+ cells, and replating. **b**) DSA induced organoids expanded for an additional 7 days co-express TP63 (green), EGFR (pink) and F3 (white). Scale bar represents 50 µm. **c**) EGFR/F3 and control FACS plots for 1 representative biological replicate (n=3 biological replicates). Isotype controls showed negative staining in 99.30%. EGFR-APC and F3-PE sorting segregated 25.90% of all cells to the double positive group. **d-e**) Immediately after cell sorting cells were cytospun on to a glass slide and stained for TP63 (red, nuclear) to identify the percentage of cells in each sorted group that were TP63+. 92.09% of double positive (EGFR+/F3+) cells exhibited nuclear TP63 staining, compared to 36.39% and 42.17% of EGFR-only and F3-only positive groups, respectively, and 8.09% of double negative (EGFR-/F3-) cells. Error bars represent the mean +/− the standard error of the mean. **f**) GFP (green) and brightfield images of whole organoids 53 days after re-plating EGFR/F3 sorted cells. Organoids were either entirely GFP+ or entirely GFP-, suggesting clonal expansion from a single cell. **g**) Protein staining of organoids derived from EGFR/F3 sorted cells revealed the presence of SCGB1A1 (pink) club-like cells and AcTUB+ and FOXJ1+ multiciliated cells (white), first two panels from left. KRT8 (green) stains the pseudo-stratified epithelium. Staining for TP63 was detected some epithelial cells (pink, right panel), and MUC5AC+ (pink) marked goblet-like cells. Scale bars represents 50 µm. **h**) Schematic of proposed (black line), hypothesized (dashed line), and inferred from previous studies (grey line) lineage relationships during human lung development.

**Extended Data Figure 10.**
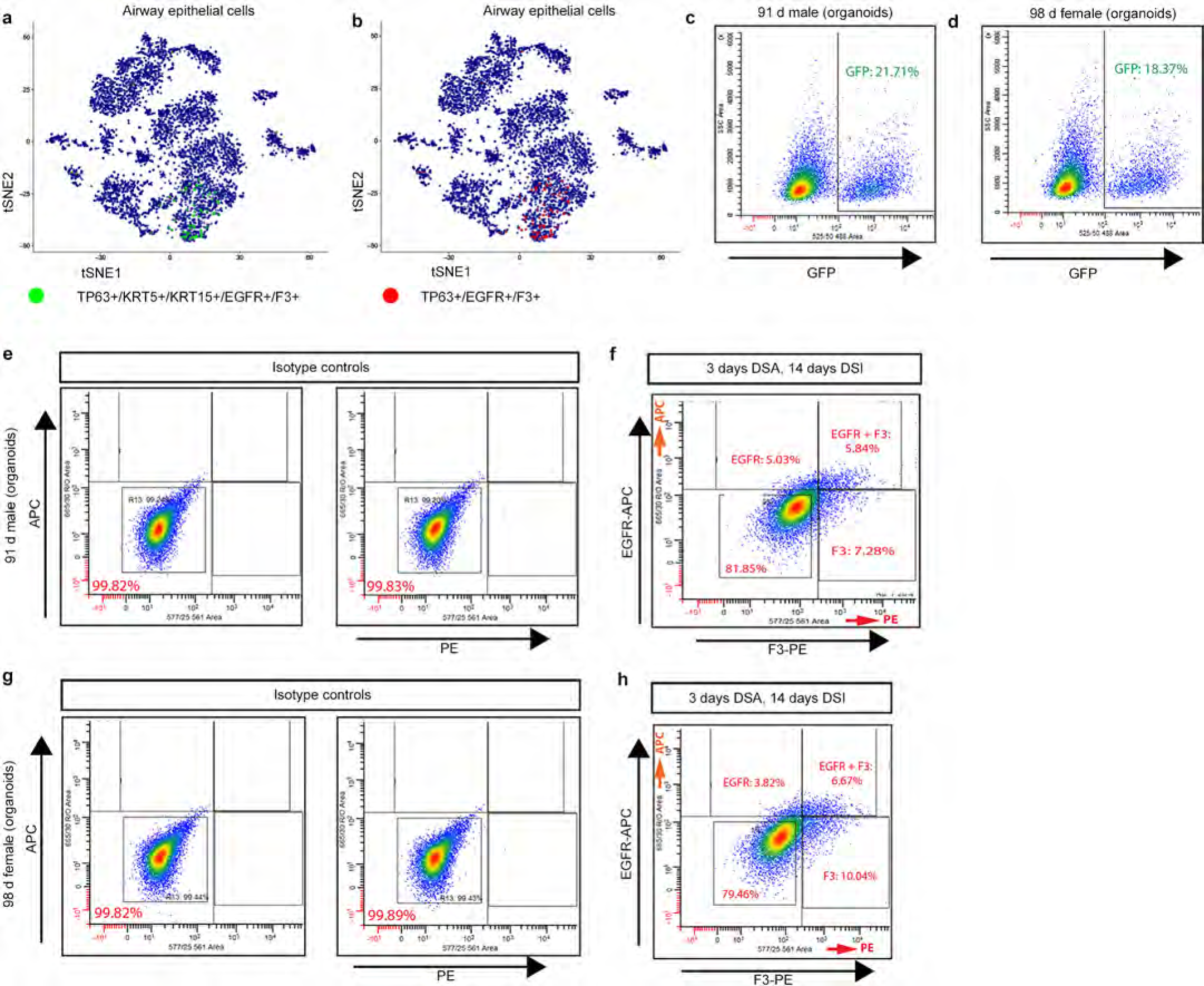
Fluorescence Activated Cell Sorting of fetal bud tip progenitor organoid-derived basal-like cells. **a**) Feature plots of all tracheal epithelial cells combined for 15, 18 and 21 week fetal lung scRNAseq from Figure 1 showing that the majority of TP63+ cells that express basal cell markers KRT5 and KRT15 and EGFR and F3 (green dots) also express TP63, EGFR and F3 (**b**) red dots). **c**) Flow sorting identified 21.71% of cells from biological replicate 2 were GFP+ and **d**) 18.37% of cells from biological replicate 3 were GFP+. **e**) Fluorescence Activated Cell Sorting (FACS) plots for biological replicate 2. Isotype controls for APC and PE show 99.82% of cells were negative for APC and 99.83% were negative for PE. **f**) Sorting on EGFR-APC and F3-PE sorted 5.84% of all cells to the double positive group. **g**) Fluorescence Activated Cell Sorting (FACS) plots for biological replicate 3. Isotype controls for APC and PE show 99.82% of cells were negative APC and 99.89 cells were negative for F3. **h**) Sorting on EGFR-APC and F3-PE sorted 6.67% of all cells to the double positive group.

This study provides a descriptive atlas of the transcriptional signatures present in different cell populations within the epithelium during human lung development. Functionally, we further identified a role for BMP/TGF/ in the transition from an undifferentiated bud tip progenitor to a proximal airway cell fate. Although future functional studies will be needed to fully define the signaling cascades that promote the transition from a bud tip progenitor to a bud tip adjacent, hub progenitor, and then basal cell, our results here demonstrate that BMP/TGFβ potently induces TP63 expression in bud tip progenitors. We hypothesize that these SMAD-induced TP63+ cells represent a very early TP63+ cell type, similar to that described at E9.5 in the primordial mouse lung bud that can give rise to all lung epithelial cell types^10^. A summary schematic of lineage relationships in human fetal lung development is proposed in Fig. 4h, with hypothesized relationships shown as dotted lines. By defining lung epithelial cell profiles throughout human development, this work serves as an important benchmark for studies aimed at differentiating lung epithelial cell types from hPSCs for drug screening, personalized medicine or regenerative medicine purposes. Similarly, the identification of SMAD signaling as a key regulator of basal cell induction may be relevant to the regulation of airway basal stem cells during injury repair across multiple organ systems, or to the appearance of basal-like populations after acute injury or in illnesses such as idiopathic pulmonary fibrosis (IPF) or basal cell-derived cancers.

## Methods

### Human Lung Tissue

Human tissue research was reviewed and approved by The University of Michigan Institutional Review Board (IRB). Human lung tissue was obtained from the University of Washington Laboratory of Developmental Biology. Tissue was shipped overnight in UWBelzer’s solution on ice.

### Paraffin processing, tissue preparation, protein staining and imaging

For all protein staining experiments, analysis was carried out on n=3 independent biological specimens, and representative images are shown in the figures. For protein analysis, tissue was immediately fixed in 4% Paraformaldehyde for 24 hours at 4°C. Tissue was washed in 3 washes of 1X PBS for a total of 2 hours, and then dehydrated by an alcohol series of each concentration diluted in 1x PBS, 30 minutes in each solution: 25% Methanol, 50% Methanol, 75% Methanol, 100% Methanol, 100% Ethanol, 70% Ethanol. Tissue was processed into paraffin blocks in an automated tissue processor (Leica ASP300) with 1 hour solution changes overnight. Paraffin blocks were sectioned with 7 µm-thick sections and immunohistochemical staining was performed as previously described^10,32^. A list of antibodies and concentrations can be found in Extended Data Table 5. All images were taken on a NIKON A1 confocal and assembled using Photoshop Creative Suite 6. Imaging parameters were kept consistent for all images of the same experiment and any post-imaging manipulations were performed equally on all images from a single experiment.

### *In situ* hybridization

For all *in situ* hybridization staining experiments, analysis was carried out on n=3 independent biological specimens, and representative images are shown in the figures. Human fetal lung tissue was fixed for 24 hours at room temperature in 10% Neutral Buffered Formalin (NBF), washed with DNAse/RNAse free water (Gibco) for 3 changes for a total of 2 hours. Tissue was dehydrated by an alcohol series diluted in DNAse/RNAse free sterile water for 30 minutes in each solution: 25% Methanol, 50% Methanol, 75% Methanol, 100% Methanol. Tissue was stored long-term in 100% Methanol at 4°C. Prior to paraffin embedding, tissue was equilibrated in 100% Ethanol, and then 70% Ethanol. Tissue was processed into paraffin blocks in an automated tissue processor (Leica ASP300) with 1 hour changes overnight. Paraffin blocks were sectioned with 7 µm-thick sections. All materials, including the microtome and blade, were sprayed with RNAse-away solution prior to use. Slides were sectioned freshly the night before the *in situ* hybridization procedure, baked for 1 hour in a 60°C dry oven, and stored overnight at room temperature in a slide box with a silicone desiccator packet, and with seams sealed using parafilm. The *in situ* hybridization protocol was performed according to the manufacturer’s instructions (ACDbio; RNAscope multiplex fluorescent manual protocol). The human *TP63* probe was generated by ACDbio targeting 4309-1404 of TP63 (accession NM_001114982.1) and is commercially available (acdbio.com, catalog number 601891-C2). The human KRT5 probe was generated by ACDbio targeting 78-2053 of KRT5 (accession NM_000424.3) and is commercially available (acdbio.com, catalog number 310241).

### Bud Tip Progenitor Organoids

All experiments were carried out using tissue from 3 different biological specimens at approximately 12 weeks gestation (91 day male, 96 day female and 98 day male). The peripheral portion of the lungs were enzymatically and mechanically disrupted to isolate bud tip epithelial cells, which were subsequently cultured in 3-dimensional Matrigel droplets with media conditions optimized to expand and maintain bud tip progenitor organoids, as previously described^11,24^. Briefly, 1 cm^2^ segments of distal lung tissue were cut from the lung with a scalpel, tissue was dissociated using lung dispase (Corning) on ice for 30 minutes, followed by incubation in 100% Fetal Bovine Serum (ThermoFisher Scientific cat. no. 16000044; FBS) for 15 minutes. Tissue was then vigorously pipetted up and down with a p200 to separate the epithelial bud tips from the mesenchyme. Tissue was washed multiple times in sterile 1x PBS to remove mesenchymal cells and epithelium-enriched bud tips were plated in a Matrigel droplet. Buds tips could be frozen down immediately after isolation or at any time after culture in 10% DMSO, 10% FBS and 80% DMEM F12.

### Culture Media, Growth Factors and Small Molecules

All experiments utilized serum-free basal medium that has been previously described^11^. Briefly, serum-free basal medium consists of DMEM F12 (ThermoFisher Scientific cat. no. 21331020 or 21331-020) supplemented with 1X N2 supplement (ThermoFisher Scientific cat. no. 17502048), 1X B27 supplement (ThermoFisher Scientific cat. no. 17504044), 1X L Glutamine (200 mM), 1X Penicilin-Streptomycin (5000 U/mL, ThermoFisher Scientific cat. no. 15140122) and 0.05% Bovine Serum Albumin (Sigma-Aldrich cat. no. A9647). On the day of use, medium is supplemented with 0.4 µM Monothio-glycerol (Sigma-Aldrich, cat. no. M6145) and 50 µg/mL Ascorbic Acid (L-Ascorbic Acid, Sigma-Aldrich cat. no. A4544, CAS Number 50-81-7). To maintain bud tip progenitor organoids in a progenitor state, serum-free basal medium was further supplemented with FGF7 (10 ng/mL, Recombinant Human Fibroblast Growth Factor 7; R&D Systems cat. no. 251-KG/CF), CHIR99021 (3 µM, Stem Cell Technologies cat. no. 72054), and All Trans Retinoic Acid (ATRA; 50 nM, Stemgent cat. no. 04-0021, CAS Number 302-79-4). Basal cell expansion medium used the same basal medium, but was supplemented with FGF10, A8301, NOGGIN and Y27632. Growth factors and small molecules were used at the following concentrations: FGF10 (500 ng/mL, made in-house as previously described^11^), A8301 (1 µM, Stem Cell Technologies cat. no. 72024), NOGGIN (100 ng/mL, R&D Systems, cat. no. 6057), Y27632 (APExBIO cat. no. A30008), LDN212854 (200 nM, R&D Systems cat. no. 6151/10), SB431542 (10 µM, Stemgent cat. no. 04-0010), TGFβ1(100 ng/mL, R&D systems cat. no. 240-B-002) and BMP4 (100 ng/mL, R&D systems cat. no. 314-BP-050).

### RNA extraction and qRT-PCR analysis

At least 1 well, containing 20-50 organoids, for each biological replicate was collected and RNA was extracted for QRT-PCR analysis. More than 1 well of organoids was collected per biological replicate and served as a technical replicate when available. mRNA for QRT-PCR was isolated using the MagMAX-96 Total RNA Isolation Kit (Life Technologies). RNA quality and concentration was determined on a Nanodrop 2000 spectrophotometer (Thermo Scientific). 100 ng of RNA for each sample was used to generate a cDNA library using the VILO cDNA kit (Invitrogen). QRT-PCR was performed on a Step One Plus Real-Time PCR System (Life technologies) using SYBR Green Master Mix (Qiagen). Expression was calculated as a change relative to GAPDH expression using arbitrary units, calculated using the following equation: [2∧(GAPDH Ct – Gene Ct)] × 10000. Some data were plotted as fold change of arbitrary expression value over a control. For this analysis, expression values for each gene for each sample, including controls, were divided by the average expression of that gene for the control group. Fold change was calculated as follows: [ExpressionGene / AverageExpressionControls] A list of QRT-PCR primers used can be found in Extended Data Table 6.

### Fluorescence Activated Cell Sorting (FACS)

To dissociate organoids into single cell suspension, organoids were first removed from the Matrigel droplet by vigorous p200 pipetting in a small petri dish filled with basal medium. Whole organoids or epithelial fragments were then transferred to a 15 mL conical tube containing 8 mL of TrypLE Express (ThermoFisher Scientific cat. no. 12605036) and placed in a tissue culture incubator at 37°C, 5% CO_2_ on a rocker for no longer than 30 minutes. Every 5 minutes, the tube was removed from the incubator and cells were agitated by pipetting the solution up and down with a P1000 pipette. Cells were confirmed to be in single cell suspension by visualization under an inverted microscope. After cells were in single cell suspension, they were spun down at 300g for 5 minutes at 4°C, resuspended in 1% BSA in HBSS, filtered through a 70 µm filter to remove any cell clusters(similar to Fisher Scientific cat. no. 087712), and spun down again at 300g for 5 minutes at 4°C. All tubes used to handle cell suspensions were pre-washed with 1% BSA in HBSS to prevent adhesion of cells to the plastic walls of tubes.

After the second wash, cells were resuspended in sorting buffer and evenly distributed in to several tubes. Cells were incubated with isotype antibody controls that were used to set FACS gates, or incubated with Anti-EGFR-APC, human (Milteny cat. no. 130-110-587, 1:50 dilution), Anti CD142 (F3)-PE, human (Milteny cat. no. 130-098-743, 1:11 dilution), REA control IgG1-APC (Milteny 130-113-434) or mouse IgG1-PE (Milteny cat. no. 130-113-762) and sorted using a Sony Synergy SY3200 system. Data was analyzed using Winlist 8.0 and FlowJo version 10.5.3 for Mac. Cells were sorted into 1mL of 1%BSA in HBSS.

### Cytospin analysis

20% of cells isolated from each group for FACS were used to evaluate the percentage of sorted cells expressing TP63. 200 µL of cell suspension (20% of 1 mL) was isolated in a separate 1.5 mL microcentrifuge tube and FBS was added to each aliquot to a final concentration of 5% vol/vol. Cells were placed in clean cytospin cones and spun at 600g for 5 minutes on a Shandon Scientific Cytospin on to charged glass slides. Slides were allowed to air dry for 5 minutes before being fixed in 100% ice cold Methanol for 10 minutes. Slides were air dried for another 10 minutes, and were then washed with 2 changes of PBS for a total of 10 minutes on a rocker. The regular staining protocol for immunofluorescence was followed (blocking, primary antibody incubation overnight at 4°C, wash, secondary antibody incubation for 1 hour at room temperature, wash, coverslip, image).

### Infection of organoids with GFP-lentivirus

Lentiviral particles were generated from a construct expressing GFP under the control of a PGK promoter with puromycin selection (Addgene plasmid #19070) by the University of Michigan Viral Vector Core. Under a dissecting microscope in a sterile hood, organoids were removed from Matrigel droplets by vigorous pipetting with a p200 pipette. Organoids were transferred to a 1.5 mL Eppendorf snap-cap tube with ∼250 µL of regular culture medium. Cells were then passaged through a 27-gauge needle attached to a 1 mL syringe 2 times in order to shear the epithelial organoids into small fragments. Cells were spun down for 5 seconds at full speed using a mini centrifuge (similar to ThermoScientific mySPIN 6, cat. no. 75004061), and the remaining floating Matrigel and culture medium were removed from the tube using the needle and syringe under a dissecting microscope. 1 mL of regular culture medium was added to the Eppendorf tube containing the organoid cell fragments, and the cells were transferred to 1 well of a 12-well tissue culture plate. 10 µM of Y27630 was added to improve cell survival and 0.5 mL of high titer virus was added to the well with cell fragments. The plate was placed in a tissue culture incubator (37°C, 0.5% CO_2_) on a rocker for 6 hours. After 6 hours, the cells suspension was moved to a 15 mL conical tube, spun down at 300g for 5 minutes at 4°C, the supernatant was removed and treated with bleach solution, and the cells were washed and spun down (300g, 5 minutes, 4°C) 3X with DMEM. On the final wash, cells were resuspended in 100% Matrigel and plated as a 3-dimensional droplet, allowed to solidify, and then overlaid with bud tip progenitor maintenance medium.

### Quantification and Statistical Analysis

All statistical analysis (quantification of immunofluorescent images and QRT-PCR data) was performed in GraphPad Prism 6 software. For quantification of protein stains in organoids, at least 3 independent organoids were counted (technical replicates) from n=3 separate biological specimens (biological replicates). All quantification of protein staining was done in a blinded fashion by an independent researcher. Statistical comparisons of data between two groups (e.g. control versus experimental condition) were made using unpaired two-sided Mann-Whitney rank-sum tests. A p value of less than 0.05 was considered significant. For QRT-PCR analysis, n=3 biological replicates were used. For each biological replicate, 1-3 well of organoids containing 20-50 organoids per well (technical replicates) was collected for analysis. To determine significance differences across multiple groups, a one-way Analysis of Variance (ANOVA) was performed followed by Tukey’s multiple comparisons analysis comparing the mean of each group to the mean of every other group. A p value of less than 0.05 was considered significant. On graphs, p values for multiple comparisons after ANOVAs are reported as follows: * p<0.05; ** p<0.01, *** p<0.001, **** p<0.0001.

### Preparation of tissue for single cell RNA sequencing

#### Human Fetal Tissue

To dissociate human fetal tissue to single cells, tissue was first dissected into regions (trachea/bronchi, small airways, distal lung) using forceps and a scalpel in a petri dish filled with ice-cold 1X HBSS (with Mg^2+^, Ca^2+^). For harvesting cells from the trachea and bronchi, the airways were transferred to a fresh petri dish filled with ice-cold 1X HBSS and were opened longitudinally with spring-loaded microscissors. The epithelium was scraped with a scalpel and a p200 pipette tip, prewashed with 1% BSA in HBSS to reduce cells from sticking, was used to collect epithelial cells and place them in a 15 mL conical tube. For distal lung tissue, a section roughly 1cm^2^ was cut with a scalpel from the most distal lung regions and minced using a scalpel and forceps. This tissue was then transferred to a 15 mL conical tube.

Dissociation enzymes and reagents from the Neural Tissue Dissociation Kit (Miltenyi, cat. no. 130-092-628) were used, and all incubation steps were carried out in a refrigerated centrifuge pre-chilled to 10°C unless otherwise stated. All tubes and pipette tips used to handle cell suspensions were pre-washed with 1% BSA in HBSS to prevent adhesion of cells to the plastic. Tissue was treated for 15 minutes at 10°C with Mix 1 and then incubated for 10 minute increments at 10°C with Mix 2 interrupted by agitation by pipetting with a P200 pipette until fully dissociated. Cells were filtered through a 70 µm filter coated with 1% BSA in 1X HBSS, spun down at 500g for 5 minutes at 10°C and resuspended in 500µl 1X HBSS (with Mg^2+^, Ca^2+^). 1 mL Red Blood Cell Lysis buffer was then added to the tube and the cell mixture was placed on a rocker for 15 minutes in the cold room (4°C). Cells were spun down (500g for 5 minutes at 10°C), and washed twice by suspension in 2 mLs of HBSS + 1% BSA followed by centrifugation. Cells were then resuspended in 1% BSA in HBSS with 0.5 units/µL of RNAseI (ThermoFisher cat. no. AM2294) in order to reduce RNA present in the media/buffer. Cells were counted using a hemocytometer (ThermoFisher), then spun down and resuspended (if necessary) to reach a concentration of 700-1000 cells/µL and kept on ice. Single cell libraries were immediately prepared on the 10x Chromium at the University of Michigan Sequencing Core facility with a target of 10,000 cells. A full, detailed protocol of tissue dissociation for single cell RNA sequencing can be found at www.jasonspencelab.com/protocols.

#### Organoids

To dissociate organoids to single cell suspensions, organoids were removed from the Matrigel droplet by vigorous pipetting with a p200 in a small petri dish, then transferred to a 15 mL conical tube containing 8 mL of TrypLE express (ThermoFisher Scientific cat. no. 12605036) and placed in a tissue culture incubator at 37°C, 5% CO_2_ on a rocker for no longer than 30 minutes. Every 5 minutes, the tube was removed from the incubator and cells were agitated by pipetting the solution up and down with a P1000 pipette. Cells were confirmed to be at single cell suspension by microscope. After most cells were in a single cell suspension, cells were spun down at 300g for 5 minutes at 4°C, resuspended in 1% BSA in HBSS, filtered through a 70 µm filter (similar to Fisher Scientific cat. no. 087712) and spun down again at 300g for 5 minutes at 4°C. All tubes used to handle cell suspensions were pre-washed with 1% BSA in HBSS to prevent adhesion of cells to the plastic walls of tubes. Cells were then resuspended in 100 µL of 1% BSA in HBSS with 0.5 units/µL of RNAseI (ThermoFisher cat. no. AM2294), counted using a hemocytometer (ThermoFisher), then spun down and resuspended (if necessary) to reach a concentration of 700-1000 cells/µL and kept on ice. Single cell libraries were immediately prepared on the 10x Chromium at the University of Michigan Sequencing Core facility with a target of 10,000 cells.

### Single Cell RNA Sequencing Computational Analysis

#### Data preprocessing and cluster identification

All single-cell RNA-sequencing was performed with an Illumina Hiseq 4000 by the University of Michigan DNA Sequencing core. Demultiplexing, initial quantification and generation of cell by gene matrices was done using 10x Genomics Cell Ranger v2.1.1-2.2.1 software with their hg19 reference. All fetal tissue samples were then combined. To ensure high data quality for further analysis, cells with more than 8,000 or less than 1,500 genes or mitochondrial transcript fraction over 10% were excluded for further analysis. Only nonmitochondrial protein coding genes were used. Gene expression levels were log-normalized by the total number of unique molecular identifier (UMI) per cell. Cellular variance of total number of UMI and mitochondrial transcript fraction was regressed out using linear regression. Highly variable genes were identified in each fetal tissue sample separately. For the combined fetal data, principle component analysis (PCA) was based on z-transformed expression levels of genes that were identified as highly variable genes in at least two samples. The top 20 PCs were used for cell cluster identification and tSNE dimensional reduction. Epithelial cell containing clusters were identified by expression of EPCAM and CDH1 and extracted for further analysis. To better resolve cellular heterogeneity of epithelial cells, sub-clustering on the extracted epithelial cells were performed. Cell clusters with low expression of EPCAM and CDH1 were removed. Based on hierarchical clustering of cluster average transcriptome and canonical cell type marker expression pattern, clusters with shared cell type identity were merged. To obtain gene signatures marking bud tip progenitors and basal cells, removal of batch effects with canonical correlation analysis, clustering and marker gene identification for fetal airway and distal lung samples were conducted for each respective group independently. Analyses mentioned in this section were mainly performed with Seurat^33^.

#### kNN network construction and visualization

Correlation distance between cells was calculated using z-transformed expression levels of genes identified as top 50 marker in any of the epithelial sub-clusters. KNN-network (k=50) was constructed. SRPING^27^ was used for network visualization.

#### Pseudotime construction

To construct the pseudotime course along the trajectory from bud tip progenitor-bud tip adjacent-hub progenitor-basal cells-differentiating basal cells, cells from the corresponding cell cluster were extracted. Based on visual inspection of kNN network visualized by SRPING, basal cell subset that do not develop into differentiating basal cells but submucosal cells were excluded. Highly variable genes were identified, and their z-transformed expression levels were used as input for diffusion map analysis using R package “destiny”^34^. Ranks of diffusion component 1 were considered as pseudotime of each cell. The pseudotime course was separated into four stages, i.e. bud tip progenitor, bud tip adjacent, hub progenitor and combined basal cell dominating stages. Cell clustering was conducted, and genes that were identified as markers of at least one clusters were considered as variable gene along the trajectory. Enrichment of BMP/TGFb signaling associated genes recorded in MSigDB^35^ among the variable genes was estimated using one-sided Fishers’ exact test.

#### Comparison of *In vitro* cell transcriptomes to *in vivo* cell transcriptomes

Data preprocessing and cluster identification of in vitro data followed the same procedure as fetal tissue data. To infer in vivo cell type for in vitro cells, correlation of gene expression levels between in vitro cells and fetal epithelial cell sub-clusters was calculated using the genes for fetal tissue kNN network construction, and assigned the in vivo cell type with highest correlation coefficients to each in vitro cell.

Detailed methods, including code used to process raw data, can be found at https://github.com/czerwinskimj/miller_lung.

### Accession numbers

Raw scRNAseq data associated with this study will be deposited in the EMBL-EBI ArrayExpress database (Accession number: **pending**). For data access ahead of final publication, contact the corresponding authors.

### Supplementary information

Supplemental information includes 10 figures, 5 tables and 2 videos.

## Acknowledgements

This work has been supported by the NIH-NHLBI R01HL119215 and Cystic Fibrosis Foundation Epithelial Stem Cell Consortium funding to JRS; AJM was supported by the Tissue Engineering and Regeneration Training Grant NIH-NIDCR T32DE007057, Cell and Molecular Biology Training Grant NIH-T-32-GM007315, and the Ruth L. Kirschstein Predoctoral Individual National Research Service Award NIH-NHLBI F31HL142197; EMH was supported by the Training Program in Basic and Translational Digestive Sciences NIH-NIDDK T32DK094775 and Cellular and Biotechnology Training Program NIH-NIGMS T32GM008353; MC was supported by the Training Program in Organogenesis Fellowship NIH-NICHD T32HD007505. The Laboratory of Developmental Biology, University of Washington, Seattle, WA, United States is supported by NIH-NICHD R24HD000836 to Ian Glass. We would like to thank Judy Opp and the University of Michigan Biomedical Research Core Facility DNA Sequencing Core. We apologize to those whose work we were unable to cite due to space limitations.

## Author contributions

AJM and JRS conceived the study. BT, GC and JRS supervised the research. AJM designed, performed and interpreted studies to characterize the native human fetal lung and *in vitro* studies utilizing fetal-derived bud tip progenitor organoids. QY, BT and GC designed and performed computational analysis on scRNAseq datasets. QY, BT, GC, AJM and JRS interpreted computational results. AW, YHT, AJM and MC collected, dissociated and submitted tissue for scRNAseq. AW and EH optimized tissue dissociation protocols for scRNAseq. MC maintained the scRNAseq database and performed quality control checks on all scRNAseq data. RC designed and performed experiments utilizing whole fetal lung explants. IG facilitated transfer of human tissue samples. AJM and TW analyzed and quantified cellular data from *in vitro* studies. YHT performed FACS experiments. AJM and JRS wrote the manuscript. QY, MC, BT, and GC provided critical feedback on the manuscript. All authors read and approved the manuscript.

## Competing interests

JRS and AJM are co-inventors on a patent filed by the Regents Of The University of Michigan relating to the isolation and maintenance of lung bud tip progenitor cells.

---

**Supplementary Table 4:** Antibody information

| Primary Antibody | Source | Catalog # | Dilution (Sections) | Clone |
| --- | --- | --- | --- | --- |
| *Biotin-Goat anti-TP63 | R&D systems | BAF1916 | 1:500 |  |
| *Biotin-Mouse anti MUC5AC | Abcam | ab79082 | 1:500 | Monoclonal |
| Goat anti-CC10 (SCGB1A1) | Santa Cruz Biotechnology | sc-9770 | 1:200 | C-20 |
| Goat anti-Chromogranin A (CHGA) | Santa Cruz Biotechnology | sc-1488 | 1:100 | C-20 |
| Goat anti-SOX2 | Santa Cruz Biotechnology | Sc-17320 | 1:200 | polyclonal |
| Mouse anti-ABCA3 | Seven Hills Bioreagents | WMAB-17G524 | 1:500 | 17-H5-24 |
| Mouse anti-Acetylated Tubulin (ACTUB) | Sigma-Aldrich | T7451 | 1:1000 | 6-11B-1 |
| Mouse anti-E-Cadherin (ECAD) | BD Transduction Laboratories | 610181 | 1:500 | 36/E-Cadherin |
| Mouse anti-F3 | Sigma-Aldrich | AMAb91236 | 1:250 | Monoclonal |
| Mouse anti-MUC5B | Abcam | AB77995 | 1:250 | monoclonal |
| Mouse anti-PLUNC | R&D systems | MAP1897 | 1:500 | monoclonal |
| Mouse anti-Surfactant Protein B (SFTPB) | Seven Hills Bioreagents | Wmab-1B9 | 1:250 | monoclonal |
| Rabbit anti-Clara Cell Secretory Protein (CCSP; SCGB1A1) | Seven Hills Bioreagents | Wrab-3950 | 1:250 | polyclonal |
| Rabbit anti-Cleaved Caspase 3 | Cell Signaling Technology | 9664 | 1:500 | Monoclonal |
| Rabbit anti-EGFR | Sigma-Aldrich | HPA018530 | 1:500 | Polyclonal |
| Rabbit anti-HOPX | Santa Cruz Biotechnology | Sc-30216 | 1:250 | polyclonal |
| Rabbit anti-IL33 | Atlas Antibodies | HPA024426 | 1:500 |  |
| Rabbit anti-KRT15 | Atlas Antibodies | HPA024554 | 1:500 | Polyclonal |
| Rabbit anti-KRT5 | Sigma-Aldrich | HPA059479 | 1:500 | Polyclonal |
| Rabbit anti-NKX2.1 | Abcam | ab76013 | 1:200 | EP1584Y |
| Rabbit anti-PDPN | Santa Cruz Biotechnology | Sc-134482 | 1:500 | polyclonal |
| Rabbit anti-phospho-SMAD1,5,8 | Millipore | AB3848 | 1:500 |  |
| Rabbit anti-phospho-SMAD2 | Abcam | AB188334 | 1:250 |  |
| Rabbit anti-Pro-Surfactant protein C (Pro-SFTPC) | Seven Hills Bioreagents | Wrab-9337 | 1:500 | polyclonal |
| Rabbit anti-SCGB3A2 | Abcam | AB181853 | 1:500 |  |
| Rabbit anti-SMAD4 | Abcam | AB40759 | 1:250 | Polyclonal |
| Rabbit anti-SOX9 | Millipore | AB5535 | 1:500 | polyclonal |
| Rabbit anti-Synaptophysin | Abcam | AB32127 | 1:500 | monoclonal |
| Rat anti-KI67 | Biolegend | 652402 | 1:100 | 16A8 |
| Secondary Antibody | Source | Catalog # | Dilution |  |
| Donkey anti-goat 488 | Jackson Immuno | 705-545-147 | 1:500 |  |
| Donkey anti-goat 647 | Jackson Immuno | 705-605-147 | 1:500 |  |
| Donkey anti-goat Cy3 | Jackson Immuno | 705-165-147 | 1:500 |  |
| Donkey anti-mouse 488 | Jackson Immuno | 715-545-150 | 1:500 |  |
| Donkey anti-mouse 647 | Jackson Immuno | 415-605-350 | 1:500 |  |
| Donkey anti-mouse Cy3 | Jackson Immuno | 715-165-150 | 1:500 |  |
| Donkey anti-rabbit 488 | Jackson Immuno | 711-545-152 | 1:500 |  |
| Donkey anti-rabbit 647 | Jackson Immuno | 711-605-152 | 1:500 |  |
| Donkey anti-rabbit Cy3 | Jackson Immuno | 711-165-102 | 1:500 |  |
| Donkey anti-goat 488 | Jackson Immuno | 705-545-147 | 1:500 |  |
| Donkey anti-goat 647 | Jackson Immuno | 705-605-147 | 1:500 |  |
| Donkey anti-goat Cy3 | Jackson Immuno | 705-165-147 | 1:500 |  |
| Donkey anti-mouse 488 | Jackson Immuno | 715-545-150 | 1:500 |  |
| Donkey anti-mouse 647 | Jackson Immuno | 415-605-350 | 1:500 |  |
| Donkey anti-mouse Cy3 | Jackson Immuno | 715-165-150 | 1:500 |  |
| Donkey anti-rabbit 488 | Jackson Immuno | 711-545-152 | 1:500 |  |
| Donkey anti-rabbit 647 | Jackson Immuno | 711-605-152 | 1:500 |  |
| Donkey anti-rabbit Cy3 | Jackson Immuno | 711-165-102 | 1:500 |  |
| Streptavidin 488 | Jackson Immuno | 016-540-084 | 1:500 |  |

**Supplementary Table 5:**
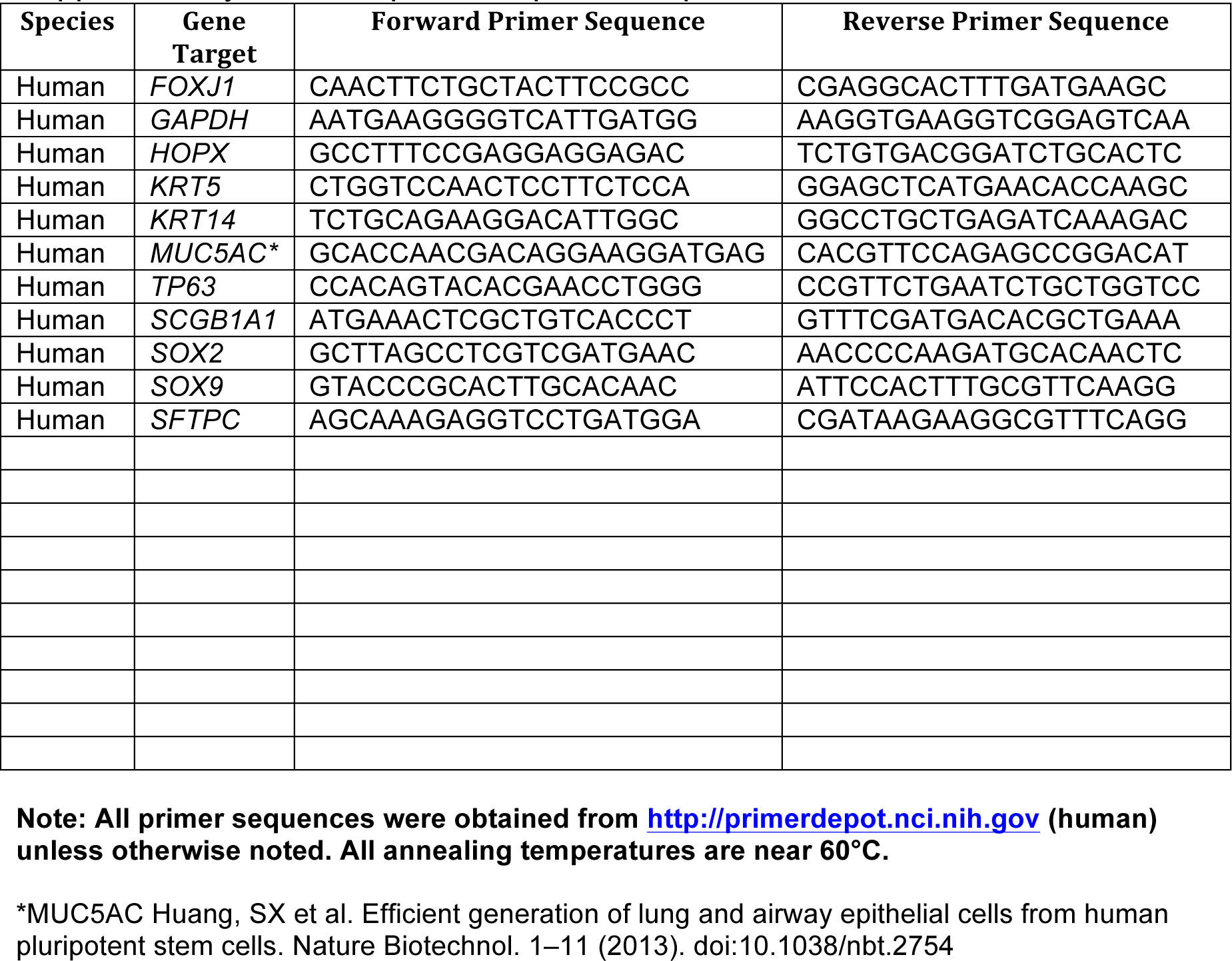
qRT-PCR primer sequences

